# High prevalence and diversity of beta-lactamase-encoding bacteria in cryosoils and ancient permafrost

**DOI:** 10.1101/2021.03.17.435775

**Authors:** Sofia Rigou, Eugène Christo-Foroux, Sébastien Santini, Artemiy Goncharov, Jens Strauss, Guido Grosse, Alexander N. Fedorov, Karine Labadie, Chantal Abergel, Jean-Michel Claverie

## Abstract

**Background:** Antimicrobial resistance is one of the major challenges affecting public health. It is mostly due to the continuous emergence of extended-spectrum β-lactamase from various environments followed by their rapid dissemination and selection in clinical settings. The warming of Earth’s climate is the other global threat facing human society, in particular with the Arctic regions experiencing a twice faster warming than the global average and permafrost affected by widespread thawing. A potentially dreadful combination of these two threats would be the release and dispersion of harmful microbes that have remained confined to largely uninhabited Arctic regions, or are stored dormant in permafrost.

**Methods:** Environmental DNA was isolated from 12 soil samples from various Arctic and subarctic pristine regions in Siberia (Yakutia and Kamchatka), including nine permafrost samples collected at various depths. The large datasets obtained from high throughput sequencing was assembled in contigs and their protein-gene contents predicted. We used exhaustive similarity searches to perform taxonomical assignments of bacterial, archaeal, and eukaryotic organisms, as well as DNA viruses. In addition, we specifically identified β-lactamase genes and their prevalence per bacterial genome estimated through the detection of two universal single copy genes.

**Findings:** A total of 9.217 10^11^ bp were exploited, leading to a total of 525,313 contigs at least 5kb in size. The DNA content of the various samples was found to be highly variable, not strictly correlated with the depth or radio-carbon-based deposit age, and most likely linked to the global density of microbes trapped in the corresponding permafrost layers. Bacteria account for more than 90% of the contigs in most samples, followed by Eukaryotes and Archaea (always lower than 10%). Viruses represented less than 2% of all contigs in all samples. The taxonomic profiles of surface cryosoils and deep permafrost samples exhibited a high diversity, including between permafrost samples originating from various depths in the same borehole. In all samples, bacterial contigs carrying different β-lactamases from class A to D were identified.

**Interpretation:** No clear common taxonomic feature could be found shared by surface cryosoils or ancient permafrost layers. However, most samples (9/12) exhibited a high frequency of β-lactamase genes, with an estimated average close to 1 copy/bacterial genome. In addition to the well-documented reactivation of infectious ancient pathogens (bacteria, viruses, protozoa), we show now that global warming could contribute to the emergence of new antibiotic resistances through the mobilization by contemporary bacteria of ancient DNA released from thawing permafrost.

**Funding:** CNRS PRC research grant (PRC1484-2018) to C.A. E C-F was supported by a PhD grant (DGA/DS/MRIS) #2017 60 0004. GG and JS were funded by ERC PETA-CARB (#338335) and the HGF Impulse and Networking Fund (ERC-0013).

## Introduction

Global warming is on the front stage of societal concerns world-wide together with the increasing frequency of emerging infectious diseases turning into pandemics (such as the latest of Covid-19)(1, 2). It turns out that both predicaments are partly linked. Through their influence on the ecology of living organisms transmitting infectious pathogens, a warming climate strongly contributes to the spread of vector-borne diseases endemic in tropical regions to more temperate regions of the globe (3–6). This phenomenon is further amplified by the encroachment into pristine areas (e.g. tropical forests) by ever expanding human activities generating new risks through imponderable contacts with (mostly uncharted) microbial environments and their associated wild life hosts (7–10).

While such dangers are mostly pointed out (particularly in Western Europe) as coming from the South, more recent concerns have been raised that new plagues could also come from the Arctic, through the release of infectious agents until now trapped in perennial frozen soils (i.e. permafrost) up to 1.5 km deep and 2-3 million years old (11–14).

Climate warming is particularly noticeable in the Arctic where average temperatures are increasing twice as fast as in temperate regions (15). One of the most visible consequences is the widespread thawing of permafrost at increasing depths (16, 17) and the rapid erosion of permafrost bluffs (e.g., 18), a phenomenon most visible in Siberia where deep continuous permafrost underlays most of the North Eastern territories.

The “ice age bug” concern is periodically brought back to the public attention. For example, when an exceptionally hot summer triggered local outbreaks of anthrax on Yamal Peninsula, Northwest Siberia, in 2016, as a deeper than usual summer season thaw of soils above the permafrost layer (i.e., the “active layer”) exhumed infectious B. anthracis endospores buried in the frozen ground since 75 years (19). Historically frequent outbreaks of anthrax killed 1.5 million reindeer in Russian North between 1897 and 1925 and human cases of the disease have occurred in thousands of settlement across the Russian North (20). Yet, anthrax is a non-contagious bacterial disease curable today given proper medical attention (antibiotics and antitoxin), and for which a vaccine is available.

Of greater concern would be the long-term preservation of animal/human viruses in permafrost. Such possibility became more realistic after the detection of variola virus DNA in a 300-year-old frozen mummy from a burial site in the Churapchinsky District of Yakutia (20, 21). However, no virus particle was likely to be infectious anymore as the viral DNA was found to be fragmented into pieces less than 2 kb (22). Such incomplete preservation could be expected given the shallow depth of the grave, at the boundary of the permafrost active layer (i.e. warming up and thawing every summers).

The capacity of deeper permafrost (hence much older) layers to preserve the integrity of “live” (i.e. infectious) viral particles, was finally demonstrated by the isolation of two previously unknown giant DNA viruses: Pithovirus (23) and Mollivirus (24). Those Acanthamoeba-infecting viruses were isolated (and cultivated) from a wall-sampled layer of a well-studied permafrost bluff (the Stanchikovskiy Yar) (Sample B, Table 1) radiocarbon dated to 35,000 years BP (23). This first demonstration that eukaryotic DNA viruses could still be infectious after staying dormant since the late Pleistocene came after many other studies (25–27) going back to 1911 (reviewed in 28). It was shown that ancient bacteria could be revived from even much older permafrost layers, up to 2 million years (29), although contaminations by modern bacteria have been suspected (30). A flower plant was also revived from 30,000 year old frozen tissues (31), as well as various fungi (from Antarctica) (32) and amoebal protozoans (33, 34).

**Table 1A:** Deep permafrost samples analyzed in this study

| Sample | IGS Code | Latitude° | Longitude° | Locality | Depth (m) | Dating Years BP | Type |
| --- | --- | --- | --- | --- | --- | --- | --- |
| P-1084T | B | 68.37016 | 161.41555 | Stanchikovskiy Yar | - | 35,000 | Wall sampling, Yedoma bluff |
| Yuk15-Yed-1 /Yed-6 | K | 61.75967 | 130.47438 | Yukechi Alas landscape | 16 | >49,000 | Core, Yedoma upland |
| Yuk15-Yed-1 /Yed-7 | P | 61.75967 | 130.47438 | Yukechi Alas landscape | 11 | >36,000 | Core, Yedoma upland |
| Yuk15-Alas-1 / Ala-11 | L | 61.76490 | 130.46503 | Yukechi Alas landscape | 12 | >28,000 | Core, from a drained thermokarst lake basin (Alas) |
| Yuk15-Alas-1 / Ala-10 | N | 61.76490 | 130.46503 | Yukechi Alas landscape | 16 | >45,000 | Core, from a drained thermokarst lake basin (Alas) |
| Yuk15-Alas-1 / Ala-9 | R | 61.76490 | 130.46503 | Yukechi Alas landscape | 19 | >42,000 | Core, from a drained thermokarst lake basin (Alas) |
| Yuk15-YuL-15/Y1 | O | 61.76086 | 130.47466 | Yukechi Alas landscape | 19 | >40,000 | Core, Yedoma upland, under lake (4 m water depth) |
| Yuk15-YuL-15/Y2 | M | 61.76086 | 130.47466 | Yukechi Alas landscape | 16 | >48,500 | Core, Yedoma upland, under lake (4 m water depth) |
| Yuk15-YuL-15/Y4 | Q | 61.76086 | 130.47466 | Yukechi Alas landscape | 6 | 53<sup>#</sup> | Core, Yedoma upland, under lake (4 m water depth) |

<sup>#</sup>: this soil sample was not frozen (talik). Data of O, M, Q: Jongejans LL, Grosse G, Ulrich M, et al. (2019) Radiocarbon ages of talik sediments of an alas lake and a yedoma lake in the Yukechi Alas, Siberia. PANGAEA, https://doi.org/10.1594/PANGAEA.904738 (41, 42).

**Table 1B:**
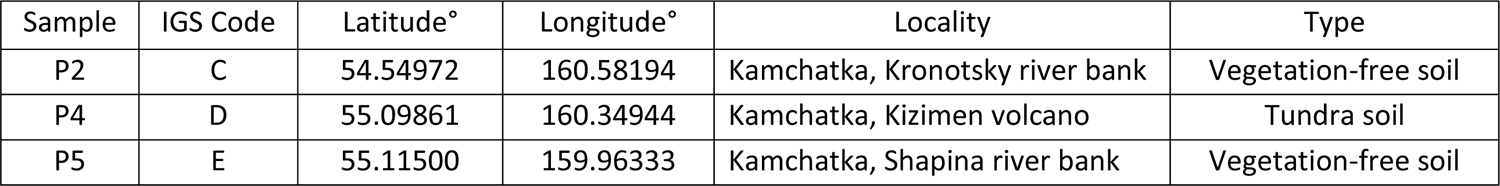
Cold surface soils samples analyzed in this study

There is now little contest that permafrost is home to a large diversity of ancient microorganisms that can potentially be revived upon thawing. Many of them, in particular among viruses, are unknown. If detected, the follow-up questions are: Are those microorganisms a threat for today’s society? Could some of these viruses and bacteria represent an infectious hazard for humans, animals or plants? Can we identify microbes the metabolisms of which would accelerate the emission of greenhouse gases (e.g. methane, carbon dioxide) upon thawing? In order to answer these questions, a first step is to document what is frozen in there. Starting this, the technique of choice (and safest) for initiating this task is metagenomics. In contrast with targeted approaches, such as metabarcoding (i.e. PCR-amplified subset of environmental DNA), the bulk sequencing of non-amplified environmental DNA allows the detection of any living organisms (and virus) without restricting the analysis to a given domain or phylum. It thus preserves the possibility of unexpected discoveries. Without amplification, it also provides an image of the real abundance of the various species, although at the cost of a lower sensitivity. Finally, the analysis of metagenomic sequences obtained from gram quantities of ancient permafrost samples is much less sensitive to eventual contaminations and more suitable for site-to-site comparisons than the use of culture-based approaches.

Here we report the metagenomics analysis of nine sub-surface Siberian samples, all taken from carbon-rich, ice-rich frozen soils called Yedoma deposits (35, 36) found in vast regions of northeast Siberia (Lena and Kolyma river basins in Yakutia)(Fig. S1) and known for well-preserved Late Pleistocene mammoth fauna remains. Yedoma permafrost may be especially prone to rapid thaw processes such as thermokarst and thermo-erosion (37) releasing not only carbon as greenhouse gases (38, 39) but also its formerly freeze-locked microbial content, in which we detected an unexpected high abundance of β-lactamases genes carried in the genomes (or plasmids) of a large phyletic diversity of cryosoil bacteria.

## Methods

### Sample collection

We sampled Yedoma and thermokarst samples from Central Yakutia from the Yukechi alas landscape about 50 km southeast of the capital of Yakutia, Yakutsk (Fig. S1). The Yukechi landscape is characterized by Yedoma uplands and drained as well as extant lake basins, indicating active permafrost degradation processes (40). Field work took place in March 2015 during a joint Russian-German drilling expedition. Four long (all ∼20m below surface/lake ice) permafrost sediment and unfrozen thaw bulb (talik) cores were retrieved. One from Yedoma deposits (61.75967°N, 130.47438°E), one below a on the Yedoma upland lake (61.76086°N, 130.47466°E), and one from the adjacent drained Yukechi alas basin (61.76490°N, 130.46503°E). Biogeochemical characterization as well as greenhouse gas production under ponding water are published in (41, 42). The Stanchikovskiy Yar sample (Sample B) from which two live viruses were previously isolated (23, 24), and three modern surface soils from pristine cold regions (Kamchatka), were also analyzed for comparison (Table 1). Fig. S1 indicates the approximate locations of the sampling sites.

### DNA extraction

Samples were processed using either the miniprep (0.25 g of starting material) or maxiprep (20 g of starting material) version of the the Dneasy PowerSoil Kit from Qiagen. Samples B, C, D, E were processed using the miniprep kit, following the manufacturer protocol and by adding DTT to the C1 buffer at a 10 mM final concentration. Extractions were repeated until 1 μg of raw DNA was obtained for each sample. All other samples (K, L, M, N, O, P, Q, R) were processed using the maxiprep kit, including the addition of DTT to the C1 buffer at a 10mM final. Samples were grounded for 20 seconds in an MP FastPrep homogenizer at a speed of 4 m/s then incubated for 30 minutes at 65 ° C and finally grounded again for 20 seconds at 4 m/s. After elution in 5 ml of elution buffer the extracted raw DNA was concentrated on a silica column from the Monarch Genomic DNA purification Kit from NEB.

### DNA Sequencing, assembly, and annotation

Each metagenomic sample was sequenced with DNA-seq paired-end protocol on Illumina HiSeq platform at the French National Sequencing Center “Genoscope” (https://jacob.cea.fr) producing datasets of 2×150bp read length, except for sample B (2×100bp read length).

Reads were quality checked with FASTQC (v0.11.7) (43). Identified contaminants were removed and remaining reads were trimmed on the 3’ end using 30 as quality threshold with BBTools (v38.07) (44). Assemblies of filtered datasets were performed using MEGAHIT (v1.1.3) (45) with the following options: --k-list 33,55,77,99,127 --min-contig-len 1000. All filtered reads were then mapped to the generated scaffolds using bowtie2 (46) with the -- very-sensitive option. To ensure a reliable taxonomic assignation, only scaffolds longer than 5kb were considered and submitted to metaGeneMark (v3.38) (47) with the default metagenome-style model. The predicted genes were then compared to the Refseq database (48) using the Diamond version of blastp (v0.9.31) (49) with an E-value threshold of 1e^-5^ and retaining only the best hit. The taxonomy of each scaffold was then inferred with a custom-made script applying a LCA like method. At each rank, recursively, the taxonomy was conserved only if at least half of the annotated genes presented the same keyword. Unclassified scaffolds and potential chimeras were screened against Refseq with Diamond blastx (--sensitive option and E-value of 1e^-5^), in order to detect exons. Non-overlapping best hits were used. The above taxonomical protocol was applied and only eukaryotic scaffolds were conserved. An alternative taxonomical annotation at the single read level was computed by Kraken (v2) (50) using its standard database. This annotation was found to be much less comprehensive and less reliable and was not used further.

### β-lactamase gene detection and analyses

ORFs with a best Diamond blastp match to a β-lactamase in the NCBI Refseq database (see above) were retrieved and analysed. Functional domains were confirmed/inferred by Interproscan (v5) (51) against Pfam, ProSitePatterns, and by Batch-CDsearch (52) against the COG database (53) (E-value < 10^-5^). ORFs at least 100 amino acids in length exhibiting a β-lactamase domain according to Pfam and COG (except for class D solely defined by COG domain) were kept for further analyses. We attempted to identify the β-lactamase-encoding contigs as potential plasmids using PlasClass (54) and PlasFlow (55). However, the results were found to be unreliable and not used further. To corroborate the high proportion of β-lactamases genes initially computed from contigs larger than 5 kb in size, the same protocol was reiterated on an independent dataset constituted of the smaller contigs in the 2kb-5kb size range. It was further confirmed by analysing a smaller open-access metagenomic dataset obtained from Arctic permafrost samples from three different locations.

Within each dataset, β-lactamase counts were normalized against three independent estimates of the number of equivalent complete bacterial genomes, using the detected number of two single copy marker genes: Phenylalanine—tRNA ligase (Uniprot accession W9B9D0) and 50S ribosomal protein L2 (NCBI accession WP_000301869.), as well as mean size of bacterial genomes. The marker proteins were first aligned to our datasets by blastP (E-value <1e-5), the resulting matching ORFs then confirmed by comparison to Refseq, and counted. The sequences of the β-lactamase-encoding contigs identified in this work will be made freely available upon publication.

## Results

### DNA content

Large variations were observed in the DNA contents of the different soil samples, despite their similar macroscopic appearances (black/brownish fine silt and sandy compact soils), including between surface samples (taken from vegetation-free spots). Using the same extraction protocol, up to 75 times more DNA could be recovered from similarly dated ancient soils from different sites (Table 1, for instance sample B *vs*. P). A lesser range of variation (10 times) was observed between surface samples (Sample C to D) already suggesting that the sample age was not the main cause of these differences. Interestingly, very similar total numbers of usable base pairs and good quality reads were determined from the 250 ng of purified DNA used for sequencing) (Table 2). Except for sample B (sequenced as shorter reads on a different platform), we observed no correlation between the initial DNA content of the sample and the number of usable good quality reads (Table 2), indicating that, once purified, all DNAs exhibited a similar quality. This suggests that the variation in DNA content is not due to its degradation, but reflects real differences in the global amount of microorganisms per gram of sample.

**Table 2:** Raw sequencing data

| Sample code | Source location | Dating (years BP) | µg DNA/ g sample | Total bp (250 ng) | Total reads (250 ng) | N contigs (>5kb) | Average size (>5kb) | Median size | Median coverage |
| --- | --- | --- | --- | --- | --- | --- | --- | --- | --- |
| C | Kamchatka | present | 10.24 | 82,686,747,816 | 676,965,994 | 42,130 | 10,314 | 7205 | 10.7 |
| D | Kamchatka | present | 0.93 | 85,127,467,969 | 642,216,312 | 78,167 | 11,551 | 7665 | 12.7 |
| E | Kamchatka | present | 1.91 | 86,068,819,231 | 685,386,422 | 16,641 | 8,659 | 6638 | 12.9 |
| B* | Stanchikovskiy Yar | 35,000 | 4.5 | 16,485,334,216 | 222,759,958 | 15,293 | 15,188 | 8583 | 13.3 |
| L | Yukechi Alas | >28,000 | 0.023 | 83,976,217,055 | 644,805,720 | 34,431 | 12,054 | 7717 | 20.1 |
| R | Yukechi Alas | >42,000 | 0.054 | 81,053,849,744 | 627,765,894 | 35,616 | 12,237 | 7787 | 19.9 |
| N | Yukechi Alas | >45,000 | 0.020 | 89,953,345,271 | 674,592,444 | 28,197 | 11,450 | 7782 | 28.1 |
| P | Yukechi Yedomas | >36,000 | 0.060 | 80,623,042,762 | 615,573,200 | 64,903 | 9,449 | 7168 | 16.3 |
| K | Yukechi Yedomas | >49,000 | 0.065 | 71,912,512,659 | 569,204,104 | 49,767 | 10,051 | 7172 | 12.9 |
| Q | Yukechi Yedomas lake | 53 | 0.067 | 82,881,074,132 | 635,044,766 | 38,175 | 15,803 | 8256 | 19.9 |
| M | Yukechi Yedomas lake | >48,500 | 0.049 | 83,875,504,131 | 648,500,972 | 58,035 | 9,764 | 7248 | 15.4 |
| O | Yukechi Yedomas lake | >40,000 | 0.051 | 77,041,387,804 | 585,962,630 | 63,958 | 9,802 | 7313 | 12.9 |

\* 100 bp/reads

In contrast, the assembly of the similar number of sequence reads from the different datasets, resulted in a highly variable number of contigs (≥5 kb). As these values are linked to the complexity of the sample (species richness-number of different species- and their relative abundance-species evenness-), this is a first indication that the samples exhibit globally different microbial population compositions and structures. This was readily confirmed by the histogram of the G+C composition of the various contigs (Fig. 1).

**Figure 1.**
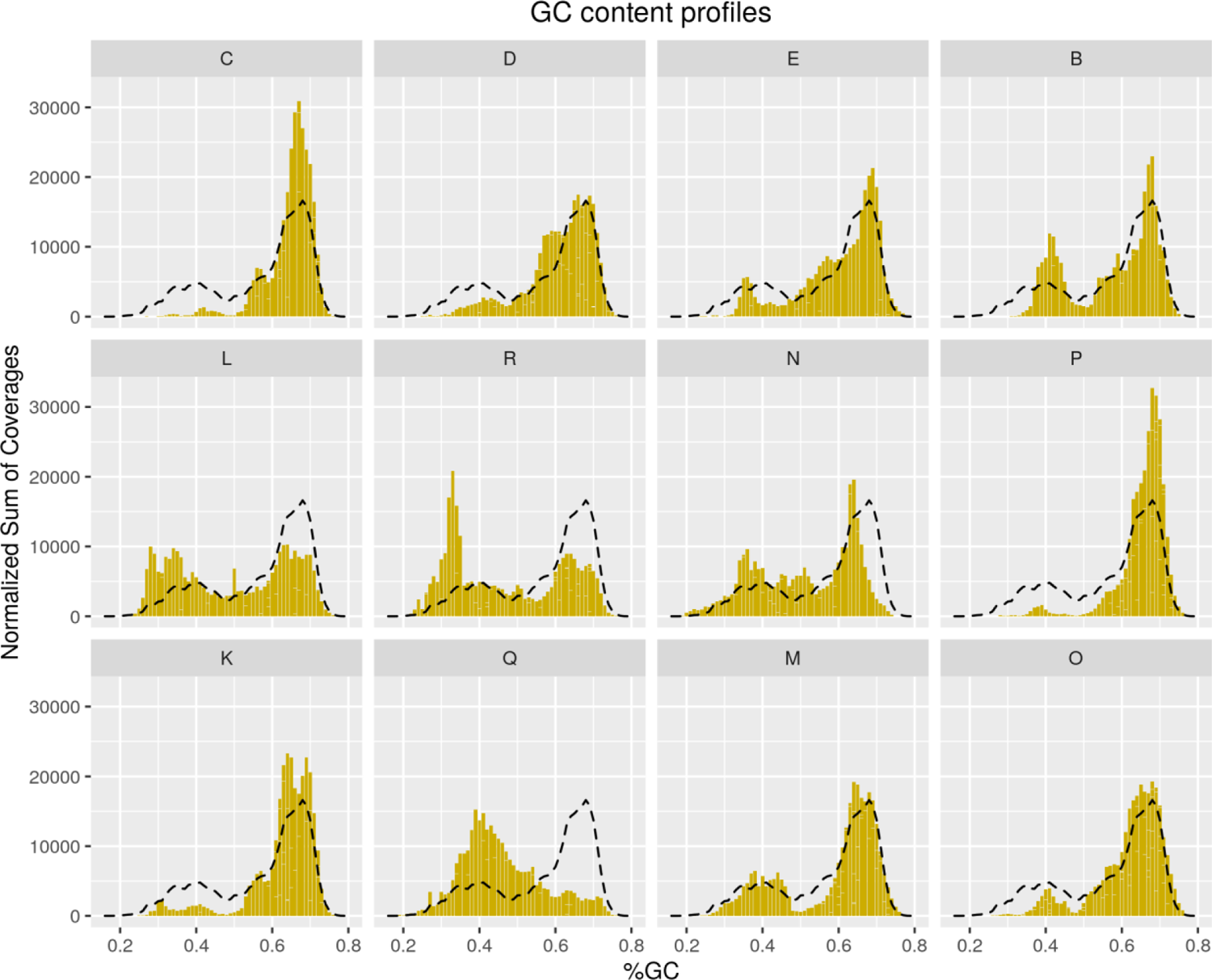
**Legend**: Global G+C content (%) in each sample, computed from their total usable bp contents. The profiles are markedly different, including for samples extracted from various levels in the same borehole: L,N,R; O,Q; K,P;. This is consistent with a minimal mixing of the microbial content. The dashed line indicates the average GC profile all samples combined.

Figure 1 illustrates the abundance of the various organisms associated with a given G+C% value in the various sample. Most of them exhibit a bimodal distribution with two peaks centered around 40% and 67% G+C, although with large variation of their relative proportions. Some of the samples exhibit almost no A+T rich organisms (e.g. C and P), while the predominant peak of G+C rich organism appears much reduced in others (e.g. L, Q, R). In two extreme cases (e.g. P and Q), a single of these peak was present suggesting the absence of whole classes of bacteria (Fig. 1). Adjacent samples from the same borehole exhibit different profiles (e.g. L, N, R and Q) supporting the absence of large-scale homogenization of the microbial populations between strata of ancient frozen soils. This validates the notion that the DNA detected at various depths truly relates to the microbes syngenetically trapped at the time of permafrost formation. The large variations of the G+C profiles between samples suggest - prior to any taxonomical annotation -, that ancient permafrost layers could harbor vastly different microbial populations. Upon thawing, the ecological or medical consequences of their eventual reactivation and release probably do not lend themselves to generalization.

### Global species contents

Compared to bar-coding methods, metagenomics allows an exploration of the sample diversity that is, in principle, only limited by a threshold of minimal natural abundance for a given species. This threshold is in fact a composite of several factors mixing the physical proportion of cells (or viral particles) of a given species (mostly unicellular) in each sample with other parameters strongly influencing our capacity to identify them by sequence similarities searches in the genomic database of known organisms. These parameters include the protein-coding density (high in viruses and prokaryotes, much lower in most eukaryotes), the fraction of the database corresponding to known members of the various phyla (high for bacteria, much lower for viruses, archaea and eukaryotes), the level of sequence similarity among homologous genes in the different domains (low in viruses, higher but variables in cellular organisms), and the proportion of ORFans in the various domains (low in Eukaryotes, high in prokaryotes, highest in viruses). Thus, for a similar coverage (i.e. relative proportion) in a metagenomics mixture, bacteria are more likely to be identified than Archaea and Eukaryotes, with viruses being the most likely to be overlooked (and remaining taxonomically unclassified). The comparisons of organism frequencies across domains are thus less reliable than within domains. Yet, the overwhelming predominance of DNA associated to the bacterial domain that characterizes all the samples is unlikely to be due to a bioinformatic/annotation bias. As depicted in Figure 2, bacteria account for at least 95% of contigs (> 5kb) in most samples. This is true of all modern cryosoils and most ancient permafrost layers despite their different origins and DNA contents. A similar predominance for bacterial was indicated using a popular read-based annotation method (Fig. S2) (50).

**Figure 2.**
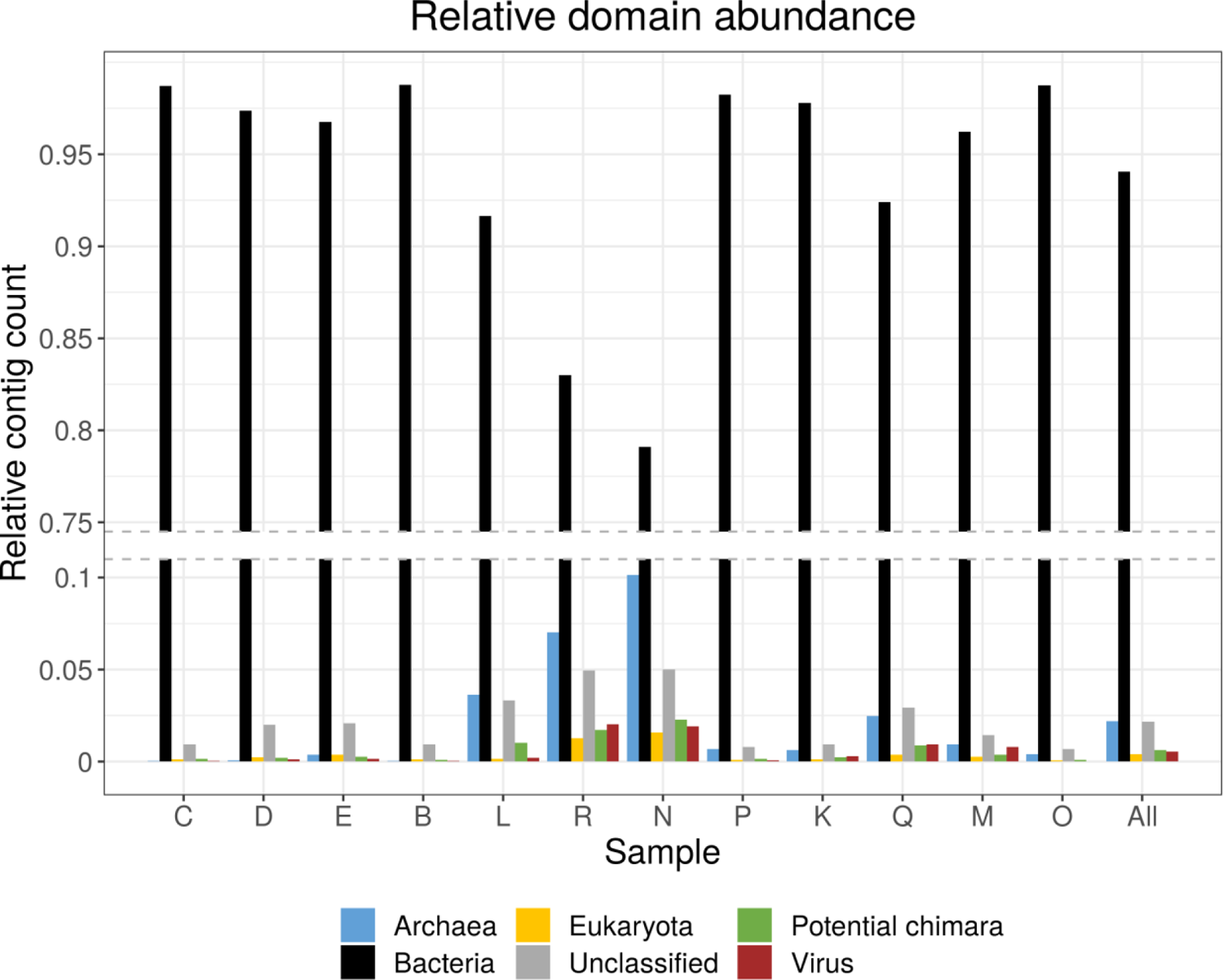
**Legend**: Relative distributions of organism types identified in the various samples. Notice that the absolute number of contigs largely varies between samples (*Table 2*).

However, we found this method to be less sensitive in identifying other cellular domains and viruses and was not used further in this study. Eukaryotic DNA is representing less than 1% of contigs, except for two sub-surface samples from the same borehole (R, N). The predicted taxonomy of these eukaryotic organisms will be described in a specific section.

A group of three samples (L, R, N) collected from the same borehole at three different depths (respectively 12 m, 16 m, 19 m) stand out from the rest by their higher proportions of contigs of archaeal origins or annotated as “chimera”. These three samples correspond to the largest median coverage values, suggesting a lower diversity of organisms (i.e. smaller richness). Contigs predicted as “chimera” probably originate from various phages the detection of which is known to be difficult in metagenomics data. A specific section will describe the types of archaea detected in the above samples.

### Focus on the bacterial populations

The relative abundances of the phyla representing more than 0.5% of the contigs are depicted in Fig.3. Even at this high taxonomical level, the picture is quite variable from one sample to the next, making it illusory to expect that a tractable number of bacterial genera could be used to classify the various types of cryosoil and permafrost into recurrent taxonomical or functional types. For instance, bacterial populations from superficial cryosoils (C, D, E) are not more similar to each other than they appear to be in ancient permafrost. Similarly, samples originating from different depths in the same borehole (such as L, R, N) do not exhibit strong similarity (ruling out by the same token the possibility of cross-contamination). Although Proteobacteria appear globally dominant (40% of the total contigs), it is not always the most abundant in every sample. The most extremes exceptions include sample P in which Actinobacteria are largely dominant, and sample Q in which Bacteroidetes is the leading phylum. The relative abundance of Firmicutes, Planctomycetes, Acidobacteria, and Verrumicrobia also vary strongly across samples from up to 24%, 18%, 13% and 8.7%, (respectively) down to almost 0. None of these characteristics appear to correlate with the geographic location or age of the ancient samples.

**Figure 3.**
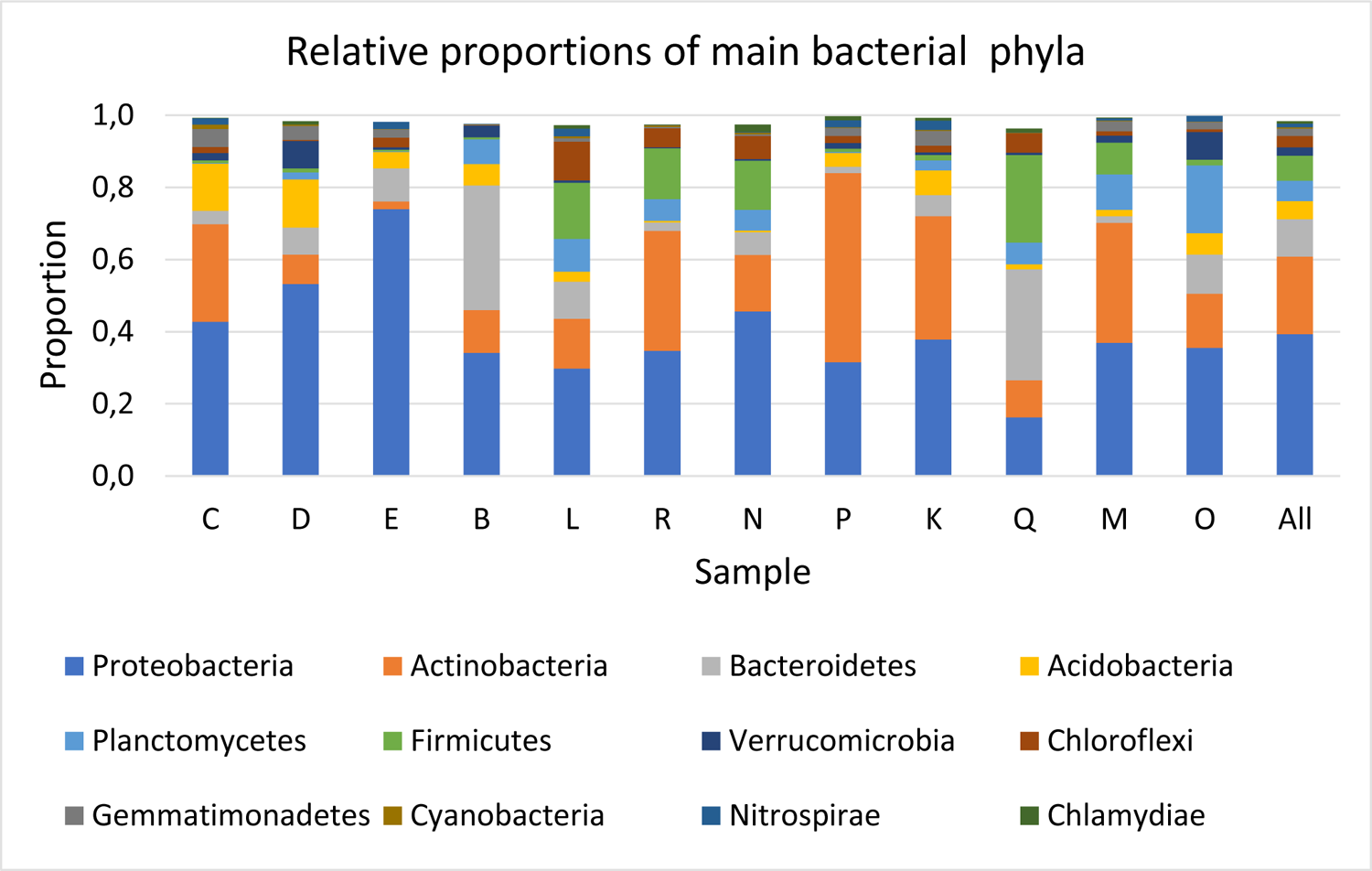
**Legend**: Relative distributions of bacterial phyla in the various sample. Only phyla representing more than 0.5% of bacterial contigs are indicated.

Even within a given phylum, such as the Proteobacteria (Fig. 4), very distinct distributions across classes could be seen between different modern cryosoils (C, D, E) or ancient permafrost layers either from the same borehole (L-R-N; Q-M-O), or from similar depth (19m: R and O). Species from the Alpha, Beta, and Delta/Epsilon divisions constitute the bulk of the proteobacteria populations, although with strong variations. For instance, Betaproteobacteria are quasi absent from the R and N samples, as are Alphaproteobacteria from sample E.

**Figure 4.**
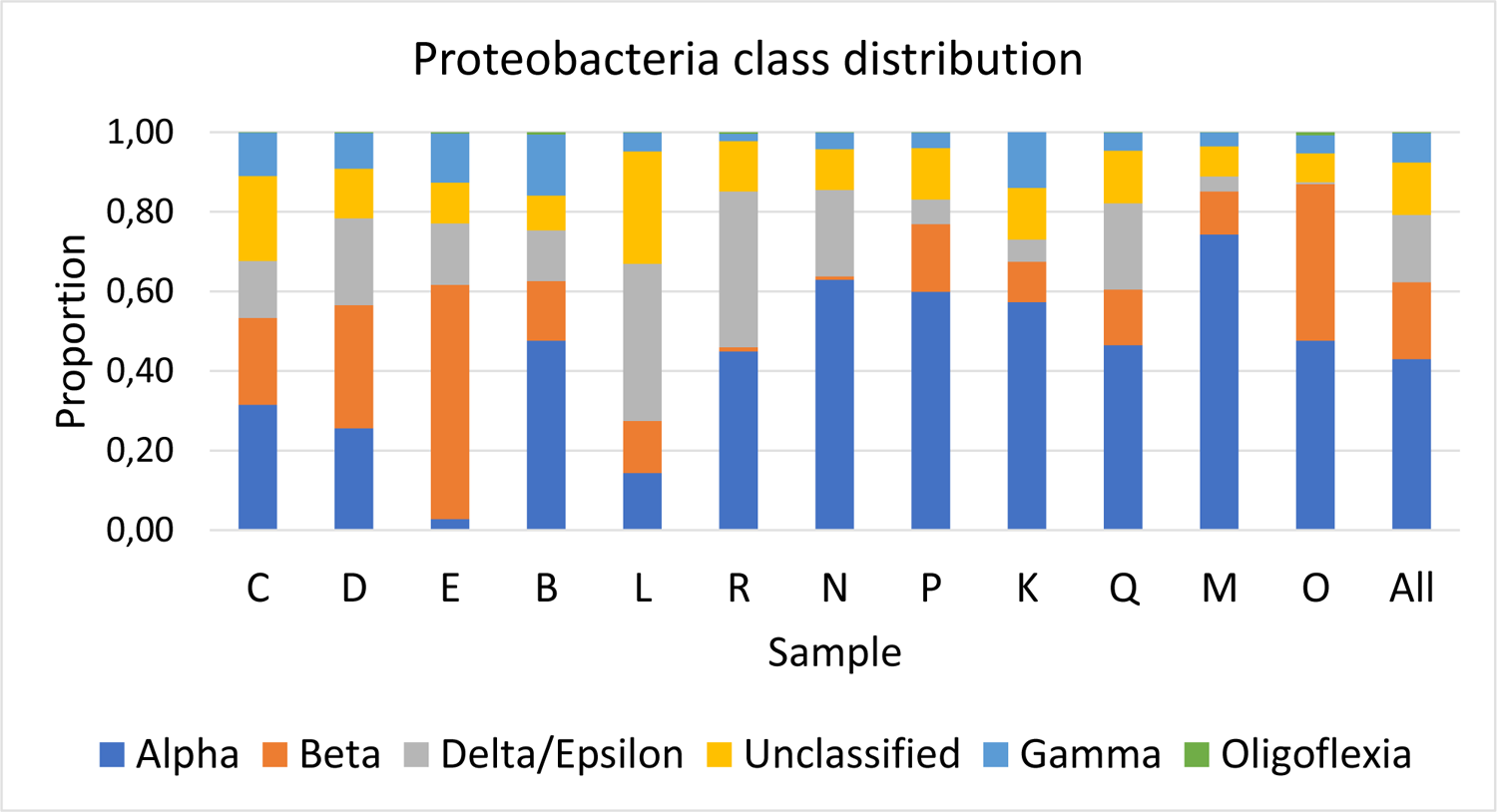
**Legend**: Relative distributions of the main clades within the dominant Proteobacteria phylum. Samples originating from various layers of the same borehole may exhibit very different profiles (e.g. L,R,N; Q,M,O), demonstrating the absence of significant mixing of their microbiome.

### Focus on the archaeal populations

The relative abundance of the main classes, orders, or phylum (at least 0.6% of the archaeal contigs) in the various samples are depicted in Fig. 5.

**Figure 5.**
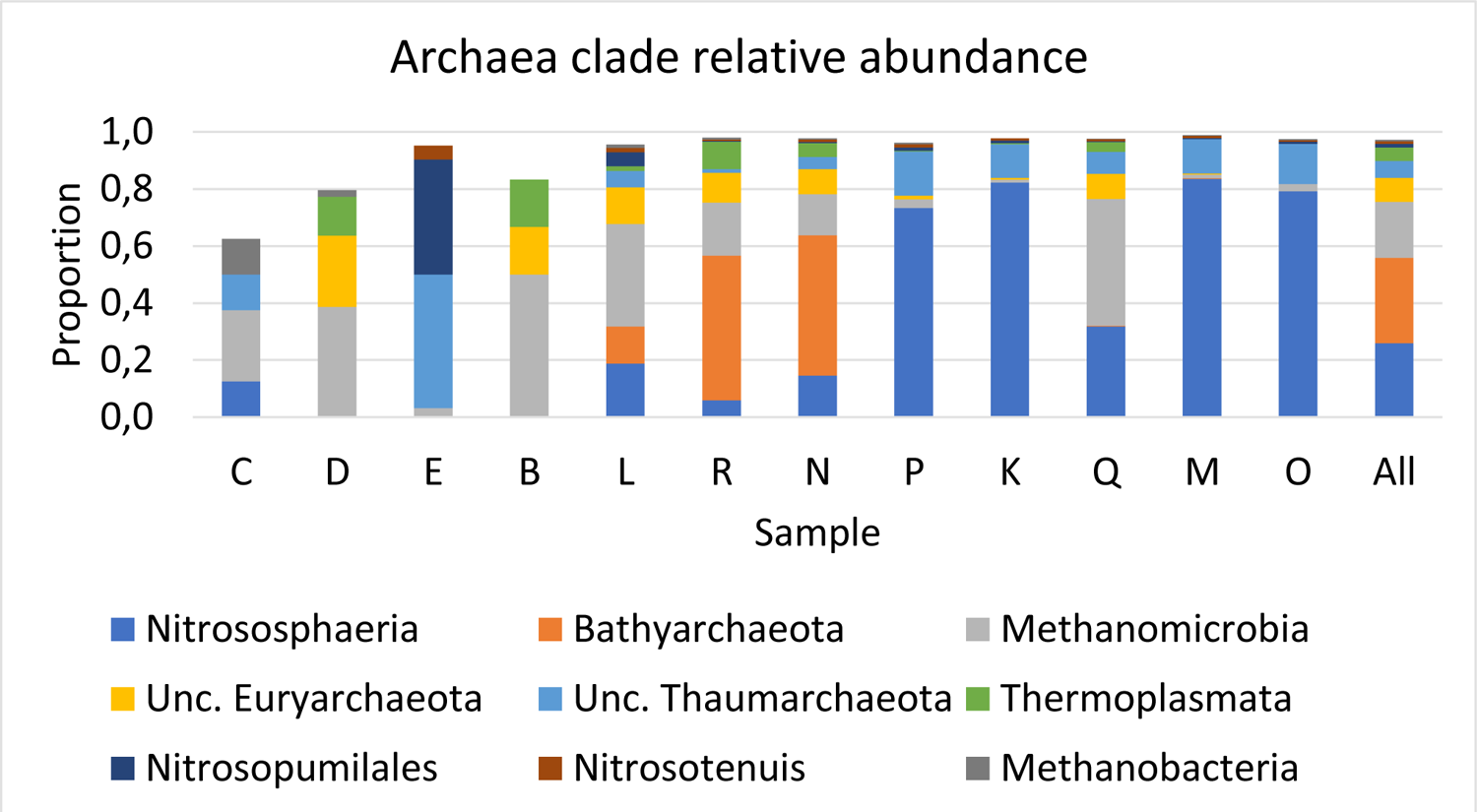
**Legend**: Relative distribution of the main clades of the Archaeal domain. The distributions are even more diverse than those computed from bacteria abundances. However, it must be noticed that the absolute number of archaeal contigs is much smaller, in particular for sample C, D, E, B (respectively 16, 44, 62 and 6 total archaeal contigs, *Fig. 2*).

This bar graph should be interpreted while keeping in mind that Archaeal contigs constitute less than 1 in thousands of all contigs in most samples (Fig. 2), except for samples L (3.5%), N (9%), R (6.75%) and Q (2.25%). As expected from anaerobic microbes, archaea are virtually absent from surface samples (less than 6/10.000 for C, D, and 4/1000 for E). In the sample where they are in sizable amounts, the archaeal population is dominated by Bathyarcheota (a recently described clade of methanogens belonging to the TACK phylum) (56), Nitrososphaeria (a class of chemolithoautotrophic ammonia oxidizing archaea belonging to Thaumarchaea phylum and exhibiting global distribution in soils) (57), and Methanomicrobia (a class of methanogens belonging to the Euryarchaeota and previously noticed to be enriched in macrophytes rhizosphere sediments) (58).

### Focus on the Eukaryote populations

The relative abundance of the main clades of Eukaryotic organisms (in term of relative number of contigs larger than 5 kb) is shown in Fig. 6. As previously (for Archaea), this bar graph should be interpreted keeping in mind that Eukaryotic contigs represent less than 1% of all contigs in most samples (Fig. 2), except for samples R (1,7%) and N (2.1%).

**Figure 6.**
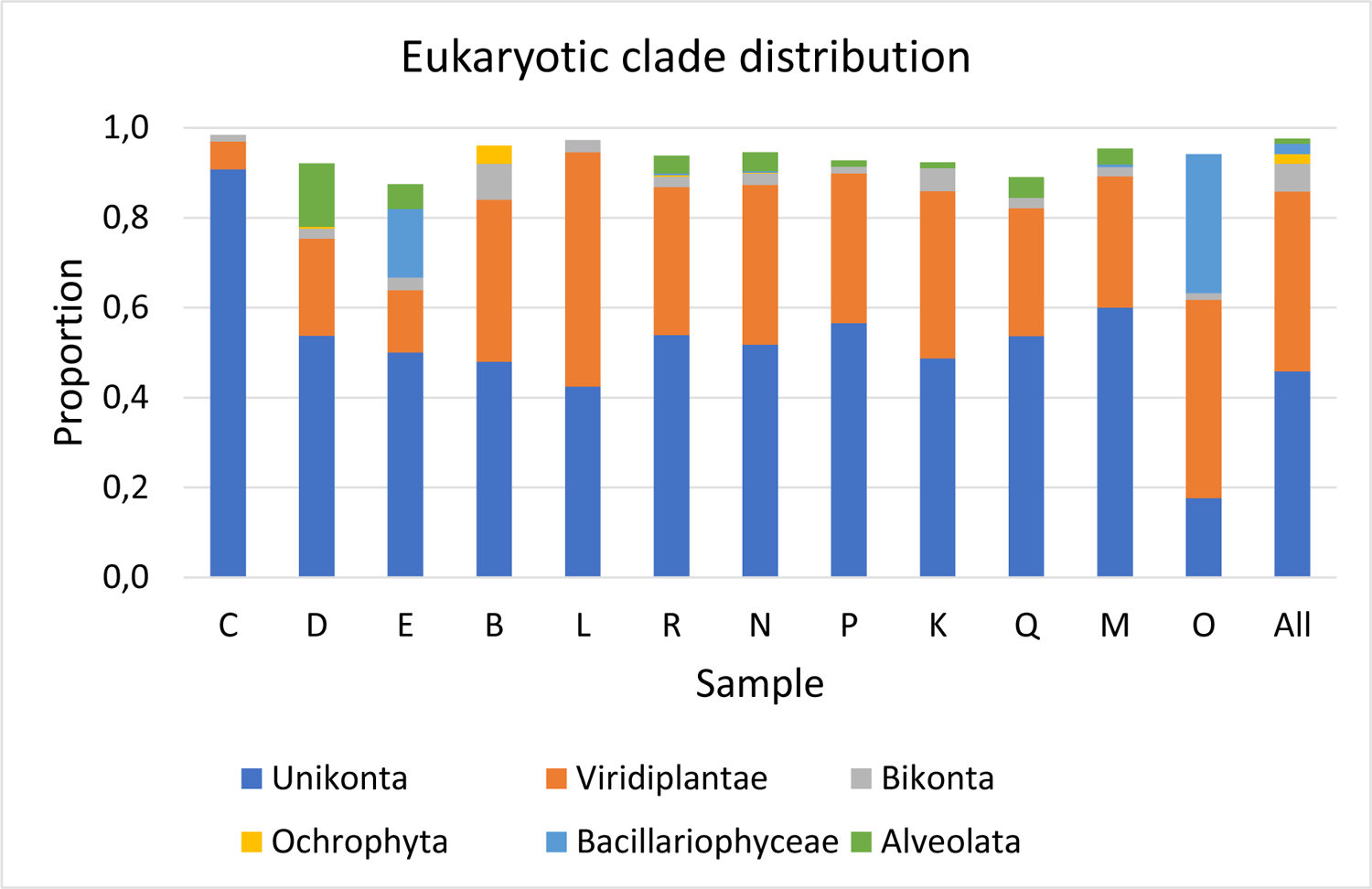
**Legend**: Relative distribution of the main clades of the Eukarya domain. It must be noticed that the absolute numbers of eukarya contigs is very small in most samples (less than 1% except for sample R and N, Fig. 2).

Despite a large variation in the proportion of Eukaryotic contigs between the different samples, the viridiplantae (land plants and green algae) and unikont (mostly fungi, metazoan and amoebozoa) constitute most of the represented clades except for sample E and O. If the later only exhibits 68 total eukaryotic contigs, 21 of which were attributed to the Bacillariophyceae (the class of Diatoms). With 30 other contigs most likely originating from green algae, this suggests that the 40,000 year-old layer corresponding to this sample was located at the bottom of a lake (Table 1A).

Concerning contigs associated to unikont organisms, the three main categories are metazoans, fungi, and amoebozoa (Fig. 7). Their relative are quite variable across different samples (although some are computed from a very small number of contigs: 12 for B and O, see Fig. 2).

**Figure 7.**
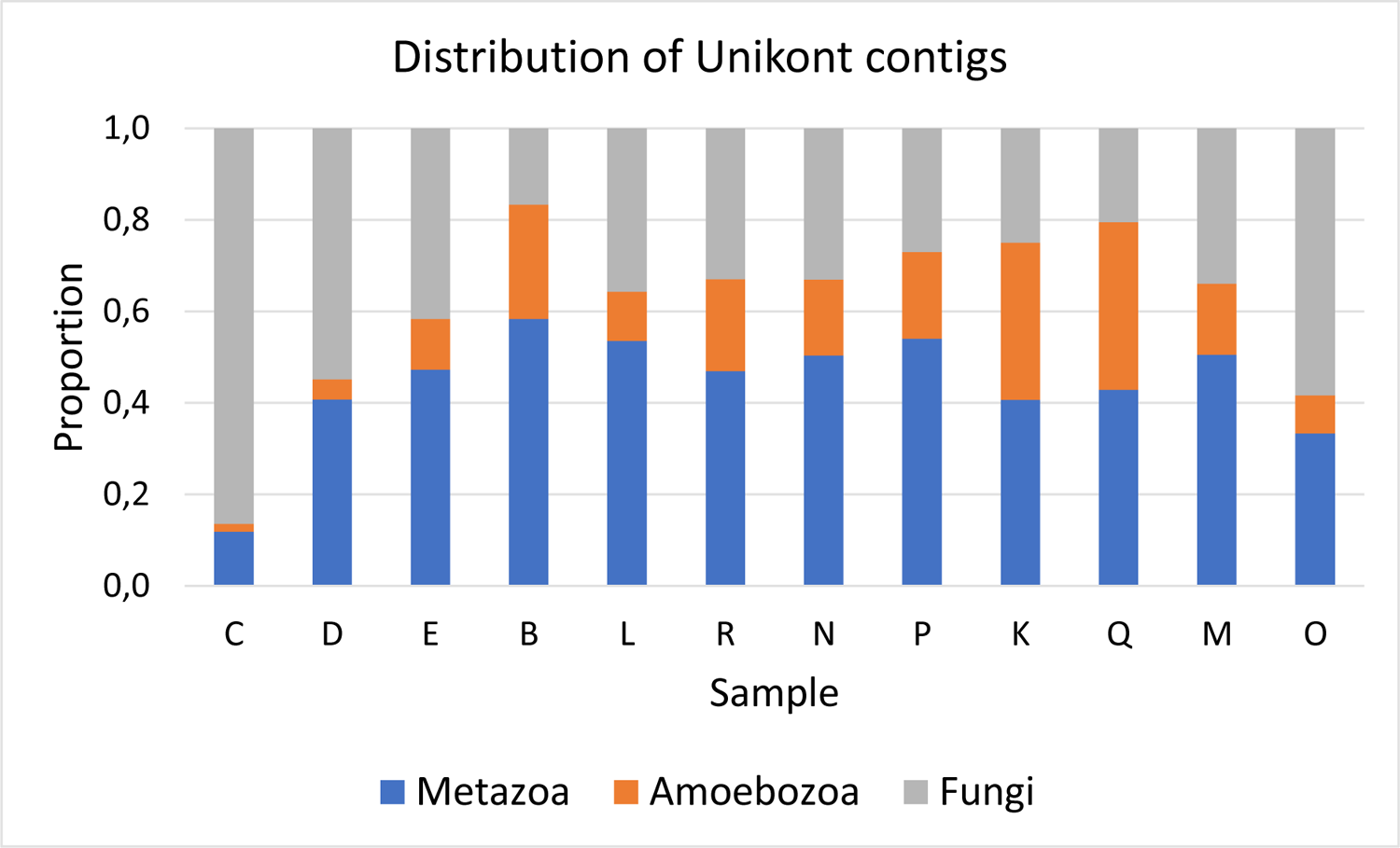
**Legend**: Relative distributions of the main clades within the Unikont phylum.

Among the samples exhibiting more than 100 unikont contigs (D, R, N, Q, M) the distributions remain quite variable, in particular between two superficial samples, both originating from Kamchatka (C, E).

### Focus on the viral populations

At a time when pandemics caused by “new” viruses seems to occur at an accelerated pace, the analysis of the viral populations present in permafrost which is known to be impacted by enhanced and deep thaw and therefore threatens to release them, is of particular interest. In particular, deep thaw by rapid thermokarst or thermo-erosion processes allows remobilization of ancient viral material tens of millennia old within short annual to decadal time scales. Unfortunately, our experimental approach is only opening a window on viruses the genomes of which consist of dsDNA, while the most human threatening viruses are RNA viruses (e.g. Coronaviruses, Influenza viruses, Ebolaviruses). These viruses are also thought to be less resistant to long period of congelation in the environment, although there is no published data on the subject to best knowledge despite the availability of potential research materials (59). We previously demonstrated the long term survival of two different types of giant DNA viruses from a sample of 28,000 year-old permafrost (23, 24) and were eager to explore how general and how diverse was the presence of DNA viruses at various locations. To our surprise, most of the samples we analyzed revealed only trace amount of viral DNA (Fig.2). Contigs classified as viral represented 0.29% of the contigs of all samples while viruses (e.g. bacteriophages) are reputed to be the most numerous microbes in soils (reviewed in 60). Viruses represented close to or more than 1% of all contigs in only 4 samples: R (2%), N (1.9%), Q (0.9%), and M (0.8%). Fig. 8 shows the proportion of main families of viruses (with more than 10 total occurrences across all samples) in the samples for which at least 40 viral contigs were detected. Although the family distributions between theses samples were quite variable, the one with the largest number of viral contigs exhibited a clear predominance of Mimiviridae, followed by Pithoviridae (the prototype of which was previously isolated from ancient permafrost (23), unclassified viruses, and a smaller proportion of phages (Caudovirales).

**Figure 8.**
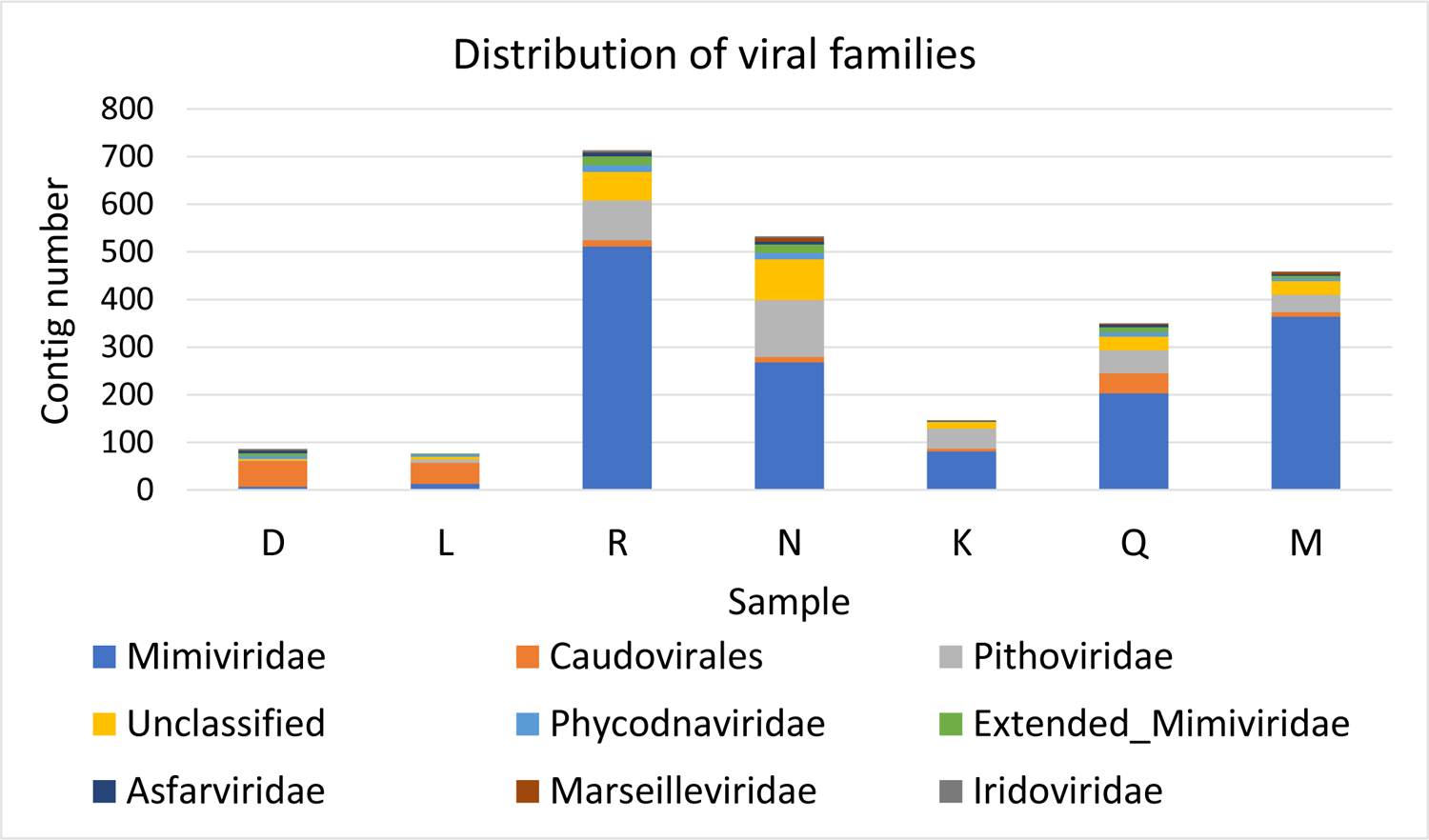
**Legend**: Relative distributions of the main DNA virus families. It must be noticed that viral contigs represent less than 1% of all contigs in most samples.

We found these viruses absent from most surface samples. Thus, we suggest that the burying of the cryosoil and its transformation into permafrost is accompanied by viral proliferations, whose genomic traces remain detectable after tens of thousands of years. As previously shown, some of these viruses can still be infectious (23, 24). The predominance of viruses of the Mimiviridae and Pithoviridae families in ancient permafrost layers correlates with the presence of their most usual cellular hosts (amoebal protozoans) in the same samples (Fig. 7). Live amoeba are routinely isolated from permafrost (33, 34). Except for a small number of contigs associated to Asfarviridae and even less to Iridoviridae, almost no eukaryotic viruses were detected from the families known to infect metazoan organisms, although their metazoan DNA was readily detected in most samples (Fig. 7).

### An unexpected large proportion of antibiotic resistance genes

Beyond the direct infectious risk represented by the re-circulation of still viable ancient microorganisms released upon thawing of old permafrost layers, the persistence of a multitude of genomic DNA fragments from all the organisms that were present in the soil at various times is also to be taken into account. Although it originated from dead (or even extinct) organisms, the permafrost DNA content constitutes a historical library of genetic resources, from which useful elements (genes) can be reintroduced into modern organisms through bacterial transformation and/or lateral gene transfer in protozoa. Of particular interest are DNA fragments coding for bacterial virulence factors, and first of all, antibiotic resistance genes the origin of which has been shown to predate by far human history (61–64). Moreover, we focused our analyses on β-lactamases the specific sequence motifs of which are readily identifiable, and for the central role the continuous emergence of extended-spectrum β-lactamases (ESBL) (64–66) play in the spread of bacterial antibiotic multiple resistances.

Fig. 9 presents the number of bacterial contigs within which β-lactamases of the 4 classes (according to Ambler’s classification) (67) were detected with high confidence (Fig. S3).

**Figure 9.**
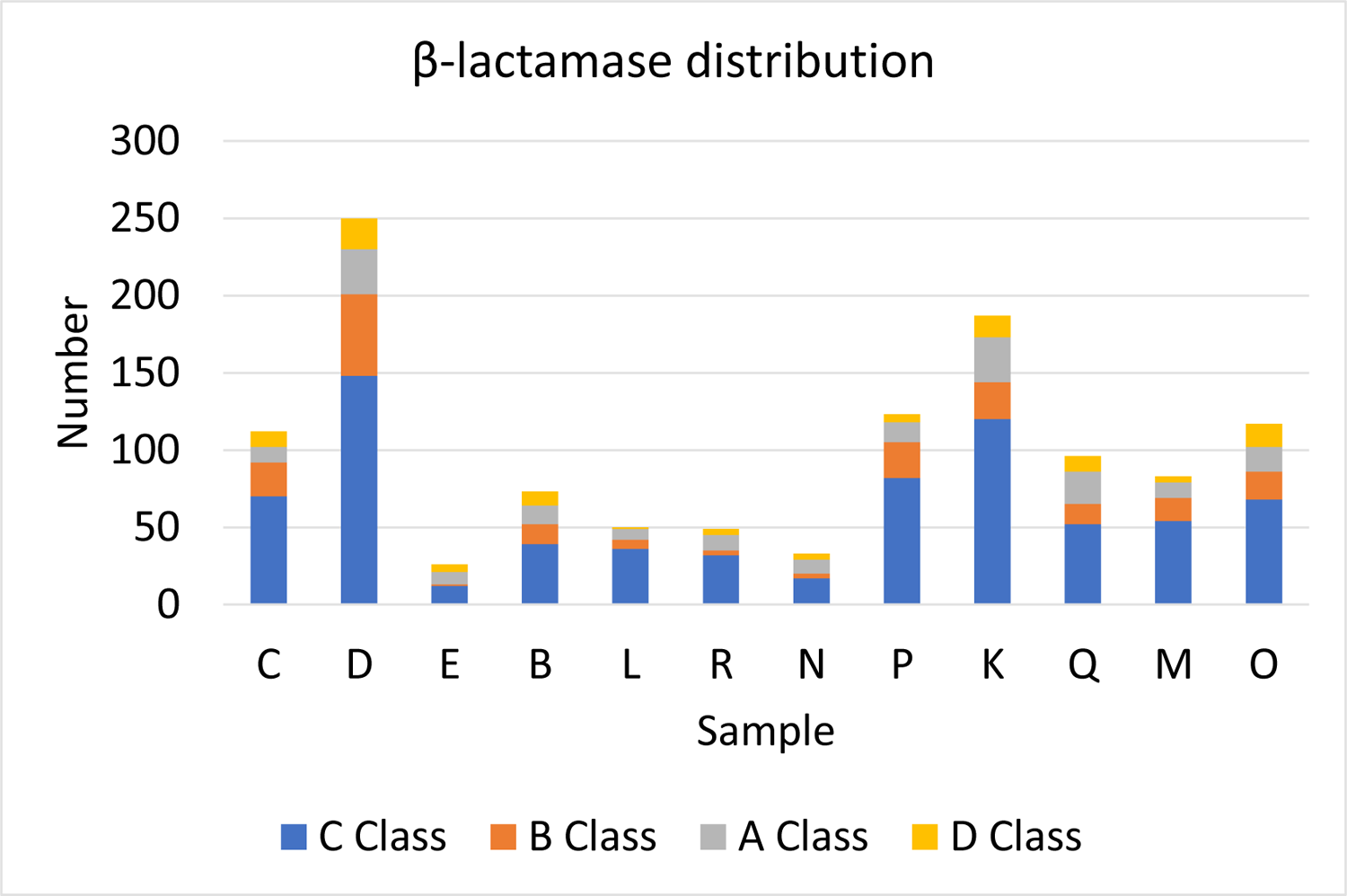
**Legend**: Relative distributions of the 4 main Ambler’s classes of β-lactamases in the various samples. See detailed statistics in Table S1.

Although these absolute numbers appear small compared to the total number of bacterial contigs (Table 2 and Fig. 2), they suggest that most of the sampled bacteria harbours close to one copy of a β-lactamase gene, once normalized by the frequency of two known single copy number genes detected in each sample data set (Fig. 10).

**Figure 10.**
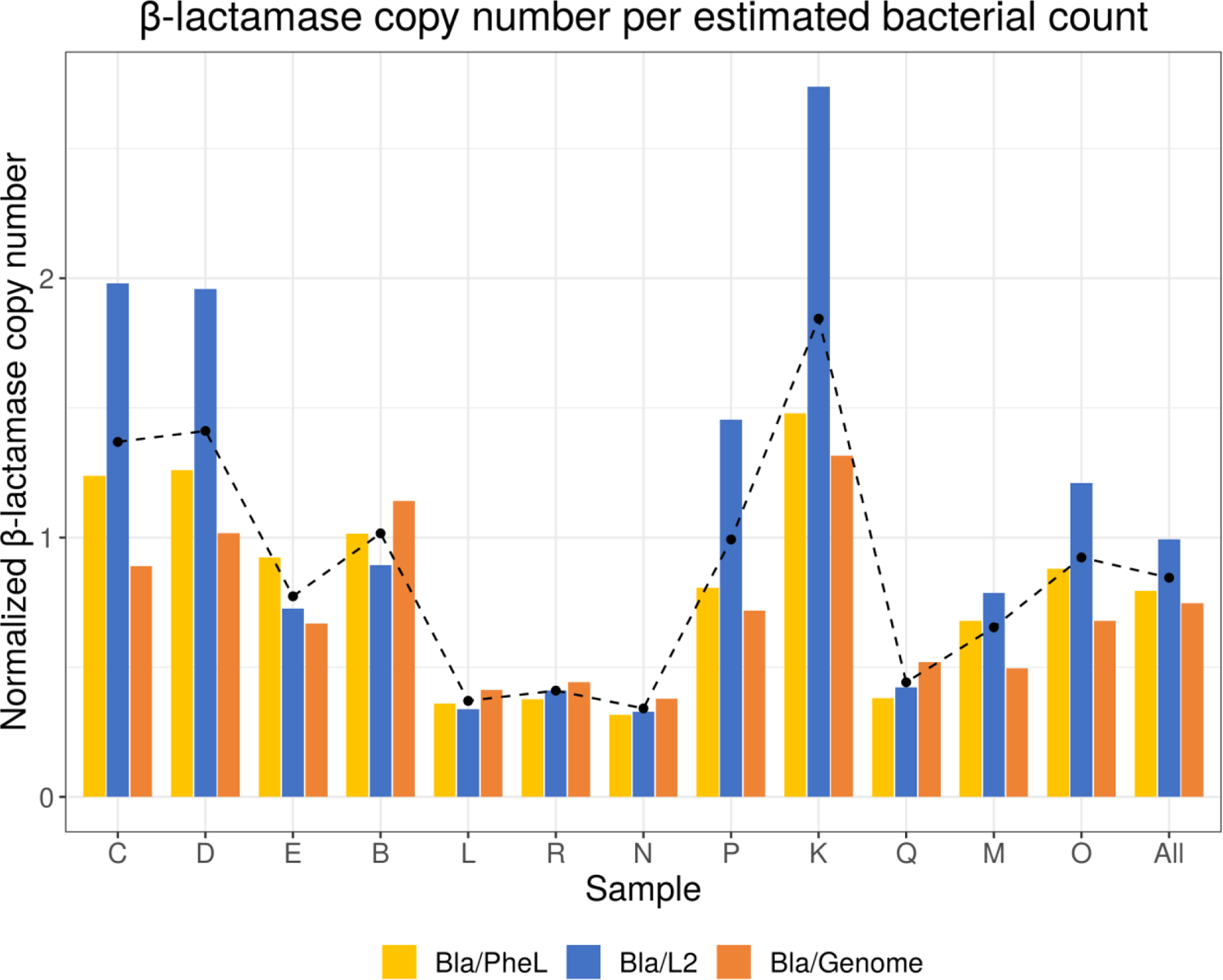
**Legend**: β-lactamase gene (Bla) copy number estimated by comparison with the abundance of two single-copy bacterial genes: phenylalanyl-tRNA ligase (PheL) (yellow), ribosomal protein L2 (blue), and the number of equivalent bacterial genomes sequenced (orange). The average of these three determination is indicated by a black dotted line. Across all samples and all references the global average β-lactamase gene copy number per bacteria is 0.87 (95% confidence interval: [0.56 – 1.17]). See detailed statistics in *Table S1*.

This copy number estimate is obtained by simply dividing the β-lactamase gene count by the number of two different universal bacterial single-copy genes, the Phenyl-alanine tRNA ligase, and the Ribosomal protein L2 (Fig. 10).

The unexpected large proportion (close to 100%) of bacteria carrying at least one β-lactamase gene indicated by the above method, was independently confirmed by calculating the ratio of the number of β-lactamases detected by the number of equivalent bacteria genomes corresponding to the total contig length for each sample. We used 3.87 Mb as the average bacterial genome sizes (68). This determination of the average β-lactamases gene copy number (0.71, 95% confidence interval: [0.51-0.]) was in good agreement with the value computed in reference to the Phenyl-alanine tRNA ligase (0.81, 95% confidence interval: [0.57-1.06], t-test p>0.45), and slightly below the one in reference to the Ribosomal protein L2 (1.11, 95% confidence interval: [0.61-1.6], t-test p>0.11).

Although a visual inspection of Fig. 10 suggests that surface cryosoils (samples C, D, E) have a higher prevalence of β-lactamase-encoding bacteria (e.g. mean: 0.87 Bla/Genome, 95% confidence interval: [0.47-1.27],) than subsurface ones (Mean = 0.66, 95% confidence interval: [0.42-0.89]), the difference is yet not statistically significant (t-test p>0.33), due to larger fluctuations in the subsurface samples. We noticed that the lowest prevalence of β-lactamase-encoding bacteria correspond to samples from the same boreholes (L, R, N, and Q, M), suggesting that it reflects an ecological reality.

Despite the above fluctuation in the average β-lactamase copy number (0.33 (N)-1.84 (K), Fig.10), the relative proportions of the 4 enzyme classes remains fairly stable across samples. Class C [52-84%] is always the most represented, followed by class B and A (respectively 16% and 14% in average), then Class D (10%) (Fig. 9).

Finally, we explored to what extent the β-lactamase genes could be unevenly distributed among the various bacterial phylum identified in the metagenomics data. For instance, one given group of bacteria could concentrate most of β-lactamase genes eventually located onto a plasmid in very high copy number. We thus simply compared the relative abundance of each bacterial phylum within the set of all cognate contigs versus those exhibiting a β-lactamase gene. Although the two distributions (Fig. 11) are far from being identical, they remain significantly correlated (Pearson coefficient > 0.8) pointing the same phyla as predominant in both distributions (Alphaproteobacteria, Bacteroidetes, Actinobacteria).

**Figure 11.**
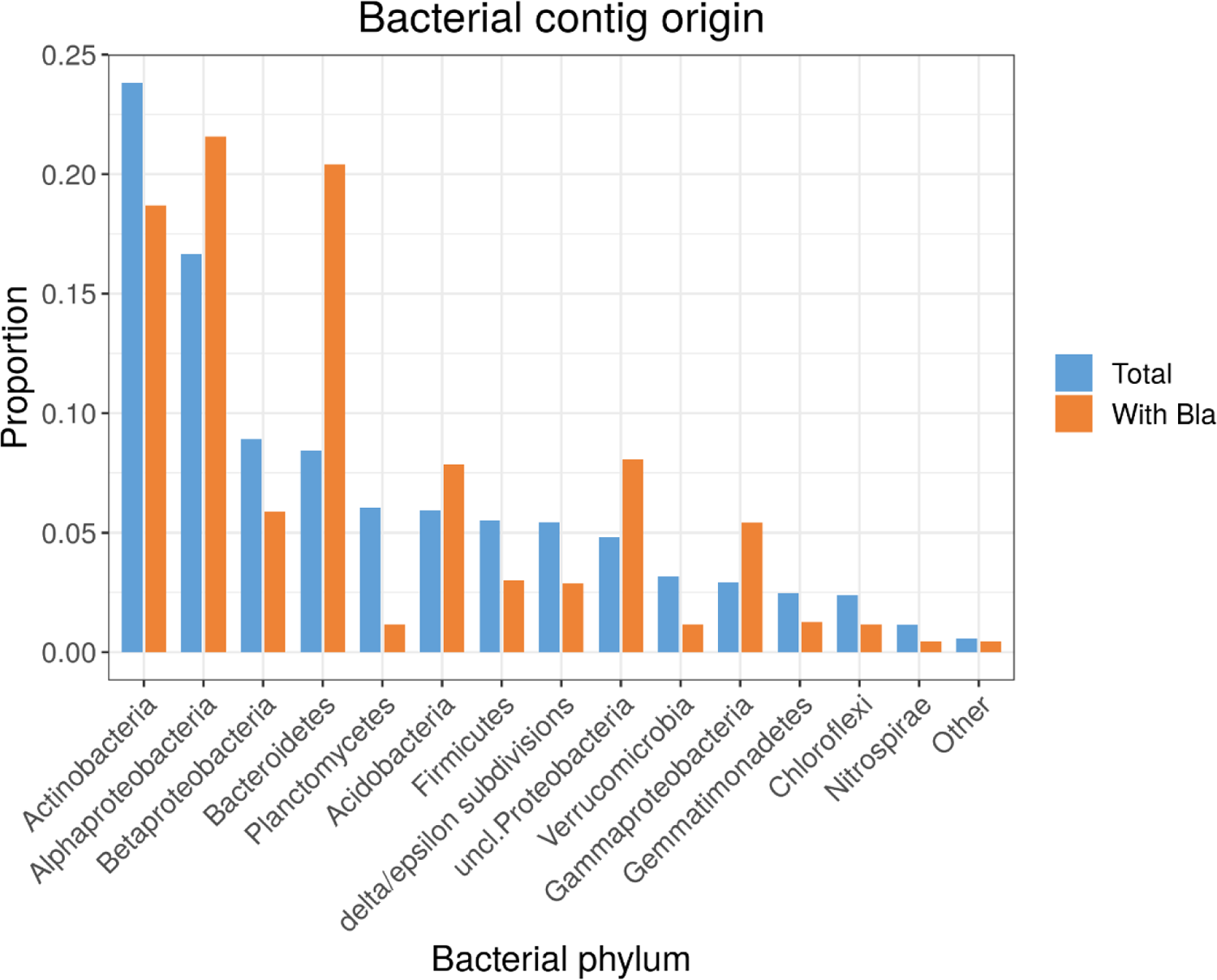
**Legend**: Side-by-side comparison of the proportions of total contigs from a given bacterial phylum with that of β-lactamase-containing genes. These proportions are globally correlated (Pearson correlation coefficient > 0.8), indicating that β-lactamases genes do not originate from a single dominant phylum. Bacteroidetes appears the most enriched phylum in β-lactamase-encoding species.

Thus, the high frequency of β-lactamase genes detected in the sampled cryosoils is not a property of a specific group of bacteria, but is shared among the very different phyla constituting these microbiomes.

We set up to further validate the above findings by two additional approaches. First we extended our analysis to contigs smaller than 5k, but still large enough to statistically encompass a complete β-lactamase coding region (about a thousand bp). The much larger number of contigs in this size range (more than 2 million, representing 1539 equivalent bacterial genomes) could thus provide an independent estimate of the proportion of β-lactamase-encoding bacteria in our sample. We summarized the results in Fig. S6. They are consistent with the β-lactamase prevalence estimated from larger contigs (mean=0.65, 95% confidence interval: [0.45-0.85]) (Table S2), although a fraction of the β-lactamase genes (about 1000 bp in size) is expected to be missed when they only partially overlap with short contigs.

In a second independent control experiment, we applied our contig assembly protocol to 3 previously published metagenomic datasets generated from deep permafrost from Yedoma deposits located on the Bykovsky Peninsula (coordinates: 72.0069 N 129.1033 E, depth: 24 m, dated >50,000 yBP), the Gydanskiy Peninsula (coordinates: 72.3741 N 78.6116 E, depth: −4.5 m, dated > 34,000 yBP), and the Omolon River mouth (coordinates: 68.7166 N 158.9000 E, depth: −26 m, dated 25,000 yBP) (69)(Table S3). These contigs were then searched for the presence of encoded β-lactamases, with the exact same bioinformatics protocol used onto our own data. Although these datasets were approximately 15 times smaller than ours (in term of total sequenced bp) and assembled in an even lesser number of contigs (Table S3) the results (although subject to larger statistical fluctuations) were again in agreement with the level of prevalence of β-lactamases computed from of our large-scale analyses (Table 3).

**Table 3:** Analysis of 3 independent datasets

| Dataset<br>Accession | site | Total bp in<br>>5kb contigs | Bla | Bact<br>genome | Bla/genome | L2 | Bla/L2 | PheL | Bla/PheL | Average |
| --- | --- | --- | --- | --- | --- | --- | --- | --- | --- | --- |
| SRX2163490 | Bykovsky P. | 33,997,197 | 4 | 8.8 | 0.45 | 10 | 0.4 | 11 | 0.36 | 0.4 |
| SRX2163491 | Omolon River | 6,278,141 | 1 | 1.6 | 0.62 | 2 | 0.5 | 2 | 0.5 | 0.54 |
| SRX2163492 | Gydanskiy P. | 61,197,557 | 8 | 15.8 | 0.51 | 6 | 1.3 | 18 | 0.61 | 0.81 |
|  | Total | 101,472,895 | 13 | 26.22 |  | 18 |  | 31 |  | 0.52 |

## Discussion

### An unexpectedly variable DNA content

We found the quantities of DNA recovered from the various cryosoils to be highly variable, from more than 10 µg/g to 0.02 µg/g, thus a factor of 500 (Table 2). Most of the low yielding samples corresponded to ancient (subsurface) permafrost layers, although without a strict correlation with age. The sample from the Stanchikovskiy Yar (B) exhibited an exceptionally high DNA contents. However, once purified, the recovered DNA exhibited similar sequencing yields (i.e. number of read/ng) and assembled into contigs of similar sizes in average (Table 2). We interpret this result as suggesting that the lower efficiency in DNA recovery from the ancient samples mainly reflects a lower global density of microorganisms rather than DNA degradation alone (at least to a point precluding its sequencing). Such a result is important because it supports the hypothesis that ancient genes carried by the ancient DNA could be recycled within contemporary microorganisms though transformation.

### Bacteria dominate DNA contents

As expected, bacteria constituted most of the DNA sources (Fig. 2). The sample displayed a large variability in the composition of their bacterial populations, as already visible from their different distribution of G+C% (Fig. 1). Such variability was reflected at the taxonomical phylum level, where the globally dominant taxa (proteobacteria, actinobacteria, bacteroidetes, acidobacteria and firmicutes) were detected in highly variable relative proportions, including among samples from different depth in the same borehole (Fig. 3).

This suggests the absence of a significant mixing between the distinct microbiomes of the permafrost layers. However, the taxonomical diversity of these microbiomes below the phylum level (and the corresponding metabolic diversity) precludes the interpretation of their composition in terms of past properties of the corresponding horizon. Such intra-phylum diversity is illustrated for the Proteobacteria in Fig. 4. There is a growing opinion that taxonomy alone is insufficient to inform us on the complexity of soil processes (70) given the already high diversity of metabolisms and life-styles encountered in a given phylum (such as the proteobacteria). Globally, the phyla found to dominate the microbiomes in our cryosoil and permafrost samples do not differ from those already reported as most abundant in previous analyses of temperate soils (71) or permafrost (72, 73). Those are proteobacteria (37%), actinobacteria (22%), bacteroidetes (12%), acidobacteria (6%), firmicutes (5%) and plantomycetes (5%) (Fig. 3). However, even the most common phyla do exhibit very different proportions between samples: 74% to 16.2% for proteobacteria, 52.4% to 2.1% for actinobacteria, 34.5% to 1.9% for bacteroidetes (Fig. 3). We noticed that the same phyla dominated both in the surface cryosoils and the permafrost sample except for the firmicutes that are almost uniquely found in subsurface samples (>1.1% at the surface, up to 24.3% in permafrost). However, this could be a bias of our limited number of samples. If not, it could be related to specific properties of these bacteria, such as endospore formation and anaerobiosis (e.g. Clostridia).

### Archeons are less abundant than expected

The production of methane, a potent greenhouse gas, is a frequently cited global-warming threat as associated with long-term thawing permafrost (42, 74, 75, and references herein). In the future, part of this flux might originate from the reactivation of methanogenic archaeons trapped in deep permafrost layers. In this respect, the very low abundance of the archaeal microbial population in most samples came as a surprise. Out of 9 analysed permafrost samples only 4 exhibited more than 1% amount of methanogenic archaeons (Bathyarcheota or Methanomicrobia)(Fig. 5). Also interesting was the fact that 3 of these samples originated from various depth (respect. 12 m, 16m and 19m) of a the same borehole. This suggests that the presence of a sizable archaeal population is not a universal property of permafrost (even within the Yukechi site), and that its detailed composition is also variable and dependent of paleoenvironmental conditions during soil formation, as previously noticed (75). Because of their apparent scarcity, the contribution of revived ancient frozen methanogens to methane production and global warming might be less than anticipated.

### A surprising scarcity of Eukaryotes

An unexpected result of our taxonomic analyses is the very low representation of eukaryotic organisms, only present in trace amounts in all our samples except for two deep samples (R, N). This result does not seem consistent with the fact that the warm spring-summer period sees the growth of a dense vegetation cover, accompanied by a variety of mosses and fungi, and a proliferation of insects, which should leave lasting traces in the metagenome of permafrost, once buried. The lack of eukaryotic contigs can be explained in several ways, in the first place by the larger size of eukaryotic genomes. For a given amount of sequenced DNA (thus a given number of cells), a larger genome will result in a smaller coverage.

However, statistically, the average size of the contigs is directly related to the coverage value. For a contig size of 5 kb, this value is ≈7, according to our assembly parameters (76). It is therefore possible that many eukaryotes do not reach such threshold. This handicap is reinforced if the proportion of eukaryotic cells is much lower than that of bacteria, as it is for archaeons. The opposite hypothesis, though less likely, could be that the eukaryotic diversity is higher, thus reducing the coverage for each of them. Fragmentation of eukaryotic coding regions by introns, and their lower density due to large intergenic regions may also strongly decrease the probability of their identification by similarity searches in protein databases.

Contigs without taxonomic assignment could therefore represent unaccounted eukaryotic contigs, but their proportion remains negligible (Fig. 2, “no taxonomy” contigs). Finally, it is also possible that the DNA of eukaryotic origin is rapidly recycled into microbial DNA during the seasonal decomposition process that takes place in the superficial active layer before being perennially frozen in the deeper permafrost.

### A low proportion of viral contigs

Contigs identified as viral origin are by far the least numerous in our analysis (Fig. 2), in particular bacteriophages (Caudovirales in Fig. 8). This is an unexpected result given that bacteria form the vast majority of microorganisms present in our samples and are generally the hosts of a wide variety of bacteriophages. One possibility is that phage populations are in fact very large, but too diverse, thus preventing the assembly of sufficiently long contigs (> 5 kb) to be formed and counted. In addition, with their very few common genes and high sequence variability, bacteriophages are notoriously difficult to identify from partial metagenomic sequences (77). They may therefore be included within contigs without taxonomic classification. Their frequent exchange of genes with different bacterial hosts can also cause them to be classified as chimeric contigs (Fig. 2). However, even considering these adjustments, their proportion would never exceed more than a few %. Finally, it is also possible that the genomes of bacteriophages, like that of other viruses, are under-represented because of the use of an inadequate DNA extraction protocol.

Regarding eukaryotic viruses, our results are nevertheless less surprising, since they show a predominance for families of viruses infecting protozoa whose presence in ancient permafrost is known (33, 34). In addition, some members of these viral families have been reactivated (23, 24). We noticed that the phylum Amoebozoa, known to be that of the hosts of families of large DNA viruses (Mimivirus, Pithovirus, Fig. 8) is among the three predominant eukaryotic clades (samples R and N, in particular).

Finally, on an optimistic note, we detected a very low proportion of viruses possibly pathogenic for animals (such as Asfarviridae and Iridoviridae), a result which puts into perspective the potential risk of viral infections triggered by the awakening of “zombie” viruses released by the thawing of permafrost. This danger could remain occasionally near the corpses of humans (21, 22) or animals (19, 20, 78, 79) well preserved in permafrost.

Nevertheless, we also have to remember that none of the viruses we managed to isolate from ancient permafrost and surface cryosoil were present in large enough abundances to achieve sizable contigs from metagenomics data (24, 80).

### Thawing permafrost as an abundant source of antibiotic resistance genes

The presence of antibiotic-resistant bacteria carrying various types of β-lactamases (A, B, C and D, according to Ambler’s classification (67)) has been previously reported in pristine Arctic and Antarctic surface cryosoils and ancient permafrost (61, 63, 81–84). However, these previous studies did not evaluate the proportion of bacteria carrying a β-lactamase gene or exhibiting a β-lactam resistance in the global population of soil bacteria, most of which are unculturable (or eventually dead, in deep permafrost).

In this work, we evaluated this proportion using two independent approaches. In the first, we compared the frequency of β-lactamase genes identified in our metagenomics data with those of two reference genes known to be present in a single copy in bacterial genomes. The second approach consisted in comparing the number of identified β-lactamase genes to the theoretical number of bacterial genomes equivalent to the total length of contig sequences. In both cases, our results indicated that more than half of the bacteria in our samples encode a β-lactamase in average (0.87, 95% confidence interval: [0.56 – 1.17]).

Similar high proportion of β-lactamase-encoding bacteria were computed from our own dataset extended to smaller contigs (Fig. S6, Table S2), and more surprisingly from metagenomics datasets generated from deep permafrost collected from remote locations (69) and different horizons. This suggests that the high prevalence β-lactamase-encoding bacteria might be a common property of cryosols, relatively independent from their detailed composition (although this would need to be confirmed by more metagenomic studies explicitly designed for this purpose).

We looked to confront our results to the existing literature. Unfortunately, we found very few studies allowing our findings to be compared with previous experimental studies. Indeed, almost all investigations of various environmental “resistomes” were performed on populations of culturable bacteria preselected for their antibiotic resistance. A few others are limited to a pre-defined bacterial species (e.g. *Escherichia coli*) (85). We could found one study analysing the proportion of β-lactam-resistant bacteria within the global bacterial population of different agricultural soils, untreated or amended with manure (86). This proportion was found to vary from 0.67% to 7.4% in untreated vs. manure-amended soils. Most of these resistant bacteria presumably encoded a β-lactamase, although to a much lesser proportion than found in our cryosoil samples.

Another shotgun metagenomics analysis of the distribution of antibiotic resistance genes (ARG) in 17 pristine Antarctic surface soils estimated the relative frequency of ARG to all genes in the [1.3-4.4 10^-5^] range which would approximately correspond (assuming 3850 as an average gene content per bacteria) to 17% of soil bacterial encoding at least one ARG (83). However, this fraction includes all ARG (e.g. multidrug resistance efflux pump, aminoglycoside acetyltransferase/nucleotidylyltransferase, aminocoumarin resistant alanyl-tRNA synthetase, etc) of which β-lactamases only constitute a small proportion.

To the best our knowledge, the high proportion of β-lactamase-encoding bacteria revealed by our metagenomics analysis of 12 diverse cryosoil samples (and 3 previously published datasets) (69), is significantly greater than what has been previously reported for other pristine environments, undisturbed by anthropogenic activities. As the function of β-lactamase in native soil microbial community is not known, our results should stimulate the building of new hypotheses. Certain soil microorganisms produce antibiotic β-lactams, including members of the Actinobacteria (e.g. Streptomyces species) (87). We noticed that Actinobacteria are among the most abundant in the cryosoil microbiomes (Fig. 3). Among eukaryotes, Fungi (also abundant in our samples, Fig. 7) have the metabolic resources to produce many complex lactam-containing compounds. They are the original source of two foundational β-lactam antibiotics: penicillin and cephalosporin (88). The higher than usual abundance β-lactamase-carrying bacteria in the cryosoils might thus reflect a particular intense war between microbes occurring during the specific perennial decomposition and freezing/thawing processes characteristic of Polar Regions. However, antibiotic resistance genes may play roles outside of the simplistic “war” paradigm. At subminimal inhibitory concentrations, antibiotics can modulate bacterial gene expression thus acting as signalling molecules rather than weapons. In such context, β-lactamases might disrupt such signalling and interfere with quorum sensing (89).

Our study indicated the presence of the four classes of β-lactamases, the evolution of which can rapidly lead to ESBL in the context of strong antibiotic selection in clinical settings. Class A EBSL includes cephalosporinases and six types of dreaded carbapenemases (90). Class B β-lactamases are metallo-enzymes using zinc at their active center compared with a serine residue for other classes. They hydrolyze carbapenems and degrade all β-lactam agents except monobactams (90). Class C β-lactamases derives from the ampC gene carried on the genome of many *Enterobacteriaceae.* Variants of these enzyme are known to reduce sensitivity to carbapenems (90). Finally, class D β-lactamases, also known as oxacillinases (OXAs) evolved from degrading an extended spectrum of cephalosporins to hydrolyze carbapenems (90). However, from their sequences alone (except for near 100% identical residues) it is impossible to predict the substrate range and the potential clinical risk of the diverse β-lactamases gene identified in our data set.

Prompted by our previous discovery of “living” viruses in Siberian permafrost (as well as several others yet to be published), we initiated this large-scale metagenomic investigation at various sites to refine our assessment of the public health risk posed by the release of unknown ancient viruses in the context of global warming and the accelerated thawing of permafrost. We found that viruses constituted the smallest part of the various cryosoil microbiomes (Fig. 2), consisting mostly of protozoan-infecting viruses and bacteriophages. Except in the immediate proximity of frozen animal or human remains, the risk of viral infections is thus probably negligible for us.

However, this study revealed a new potential danger, consisting in the unexpected large abundance of β-lactamases genes carried in the genomes (or plasmids) of a large phyletic diversity of cryosoil bacteria. The DNA from these bacteria, dead, alive, or in a frozen dormancy, thus constitutes an immense reservoir of historical antibiotic resistance genes the transfer of which to contemporary human bacterial pathogens in a clinical setting might further contribute to the antibiotic resistance crisis, now considered one of the biggest public health challenges of our time.

In a context of global warming, and in particular rapid Arctic warming, thawing permafrost will behave as a resistance gene-flavoured ice cream to be enjoyed without moderation by all bacteria passing by.

## Acknowledgements

We are deeply indebted to our volunteer collaborator, Alexander Morawitz, for collecting the Kamchatka soil samples. We thank the PACA Bioinfo platform for computing support. We thank M. Ulrich (DFG project UL426/1-1) and P. Konstantinov for helping with fieldwork at the Yukechi site as well as the Alfred Wegener Institute and Melnikov Permafrost Institute logistics for field support and sample acquisition.

## Contributions

JMC and CA designed the study, SR and SS performed the bioinformatics analyses, EC processed the sample and performed DNA extraction, KL supervised DNA sequencing, GG, JS, and ANF collected the deep core samples from the Yukechi site. All authors contributed to the manuscript writing.

---

**Supplementary Figure S1.**
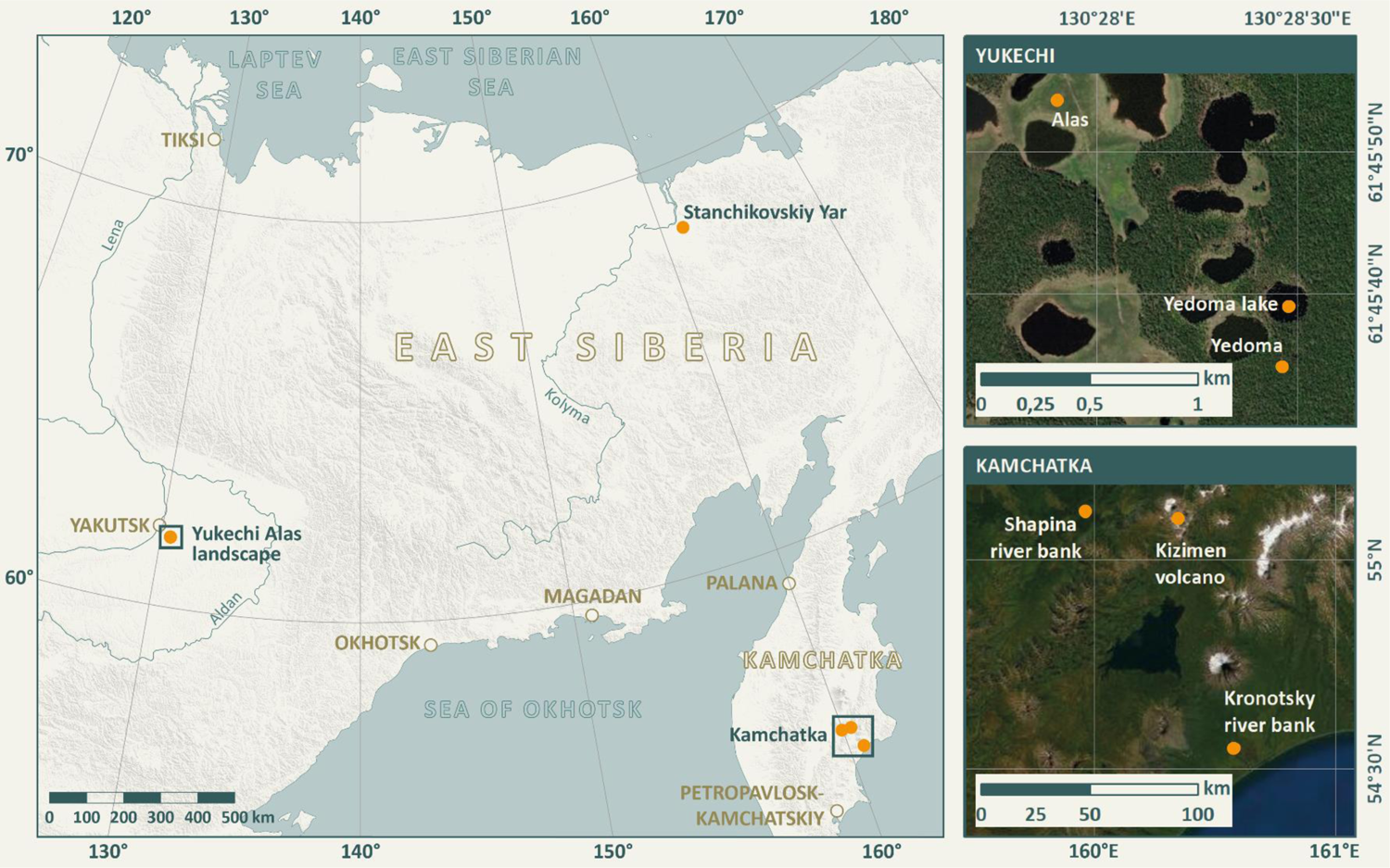
Legend. Overview of sampling locations (left map) with details on the sampling sites from Yukechi (upper right) and Kamchatka (lower right)

**Supplementary Figure S2.**
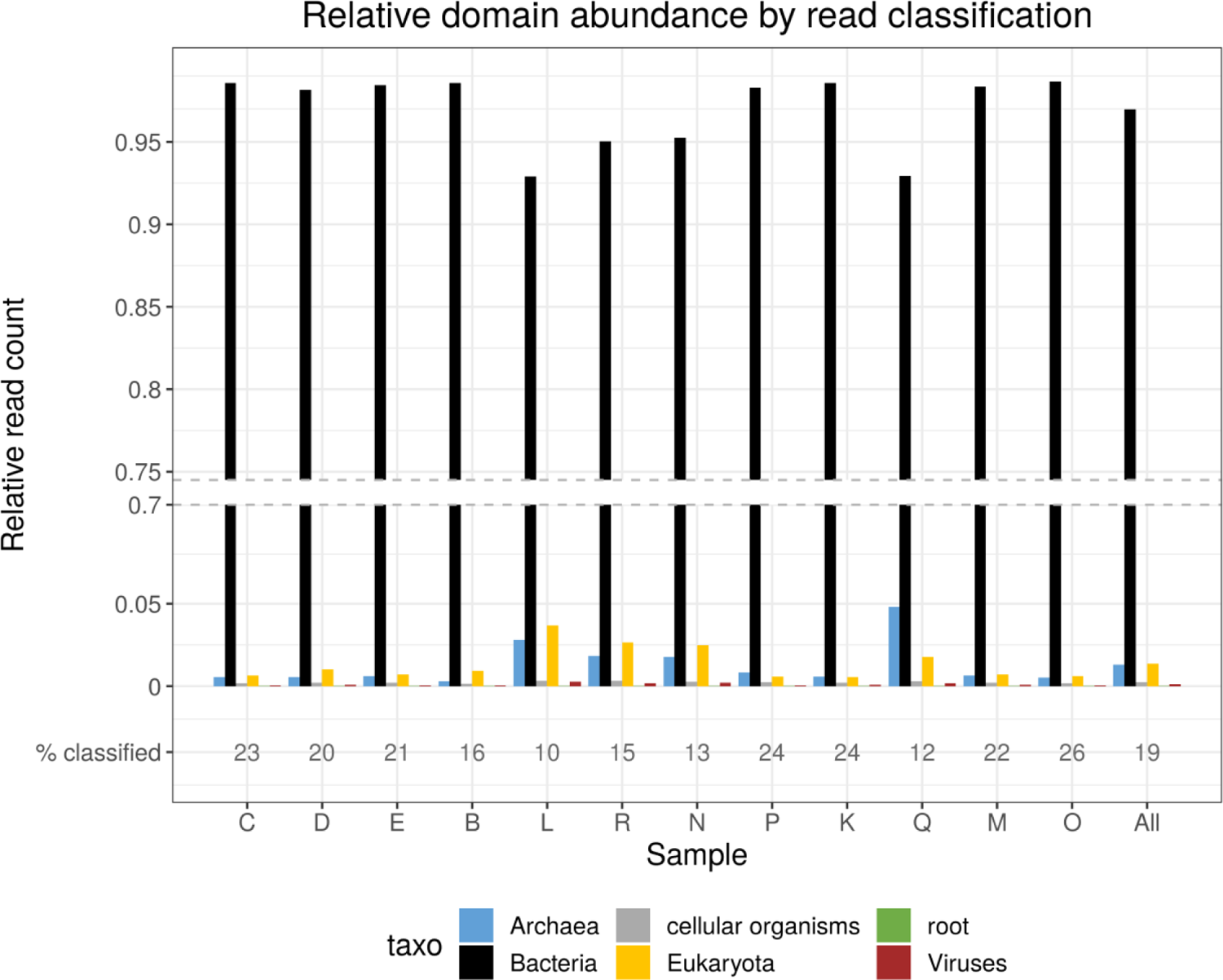
**Legend**: Relative distributions of organism types identified in the various samples, using a read-level algorithm (Kraken). Notice the small percentage (% classified) of used reads.

**Supplementary Figure S3.**
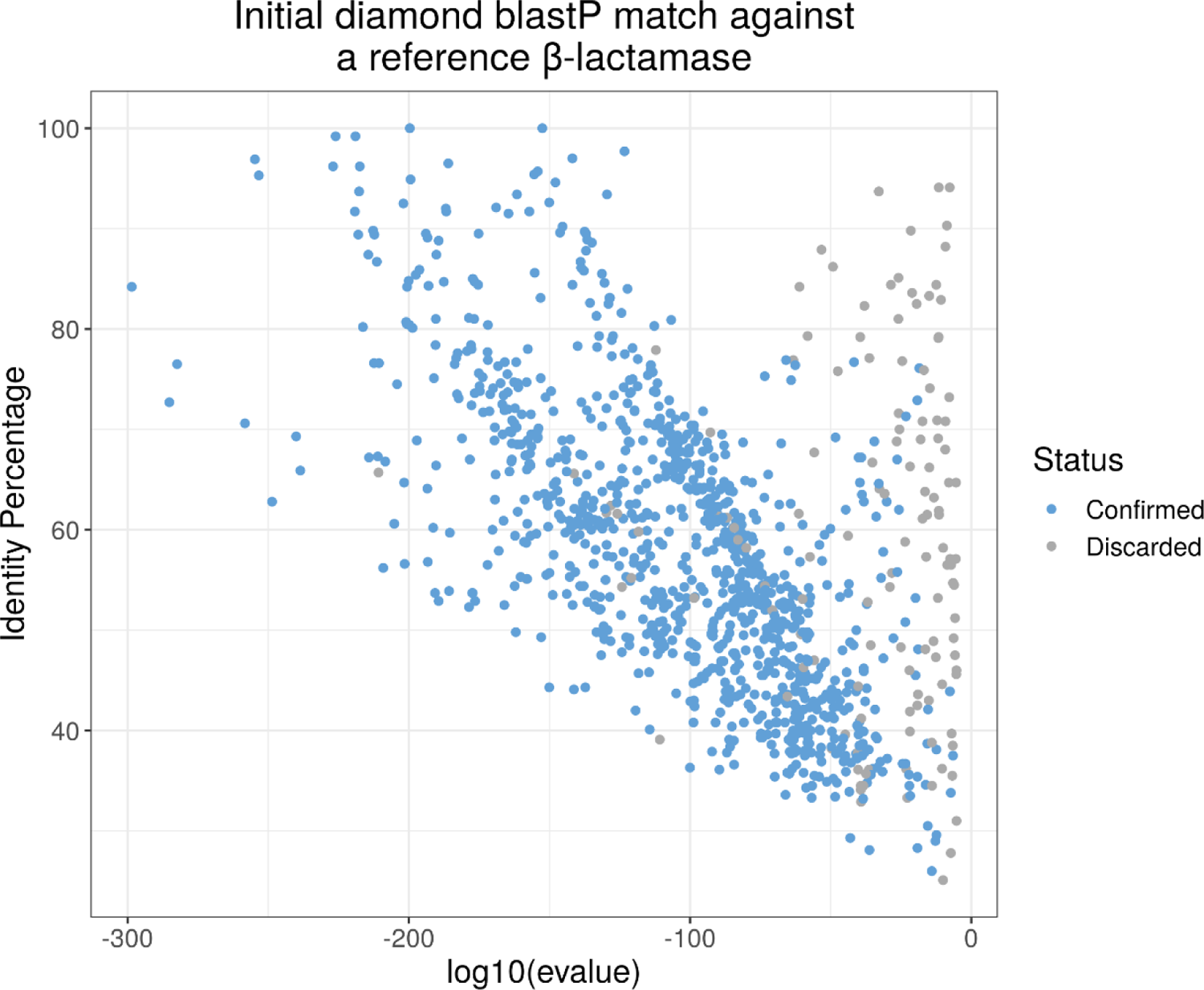
**Legend**: Similarity (% identical residues) and matching P-value of predicted ORFs with a best match with a β-lactamase in the Refseq database using Diamond. ORFs were then discarded or confirmed based on size and the presence of a *bona fide* β-lactamase domain (See Material and methods). The bulk of predicted sample β-lactamases are in the unambiguous [30%-90%] identity range.

**Supplementary Figure S4.**
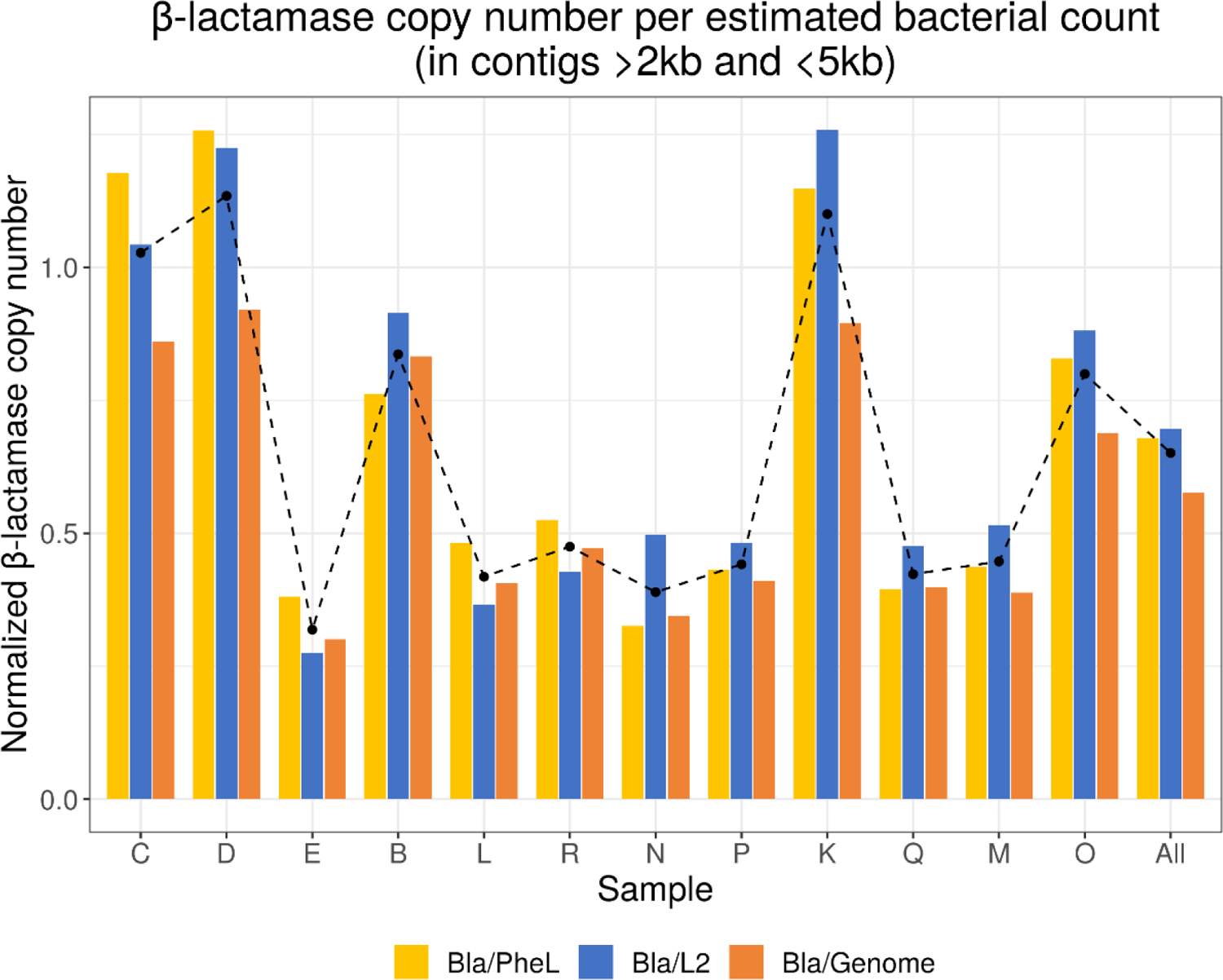
**Legend**: β-lactamase gene (Bla) copy number estimated by comparison with the abundance of two single-copy bacterial genes: phenylalanyl-tRNA ligase (PheL) (Blue), ribosomal protein L2 (orange), and the number of equivalent bacterial genomes sequenced (grey). The average of these 3 determination is indicated by a black dotted line. Across all samples, the global average β-lactamase gene copy number per bacteria is *0.65 (95% confidence interval: [0.45 – 0.84]). A very similar picture is obtained when applying a less stringent criteria not requesting each sequence to encompass the relevant complete β-lactamase domain (Fig. 10)*.

**Supplementary Table S1.**
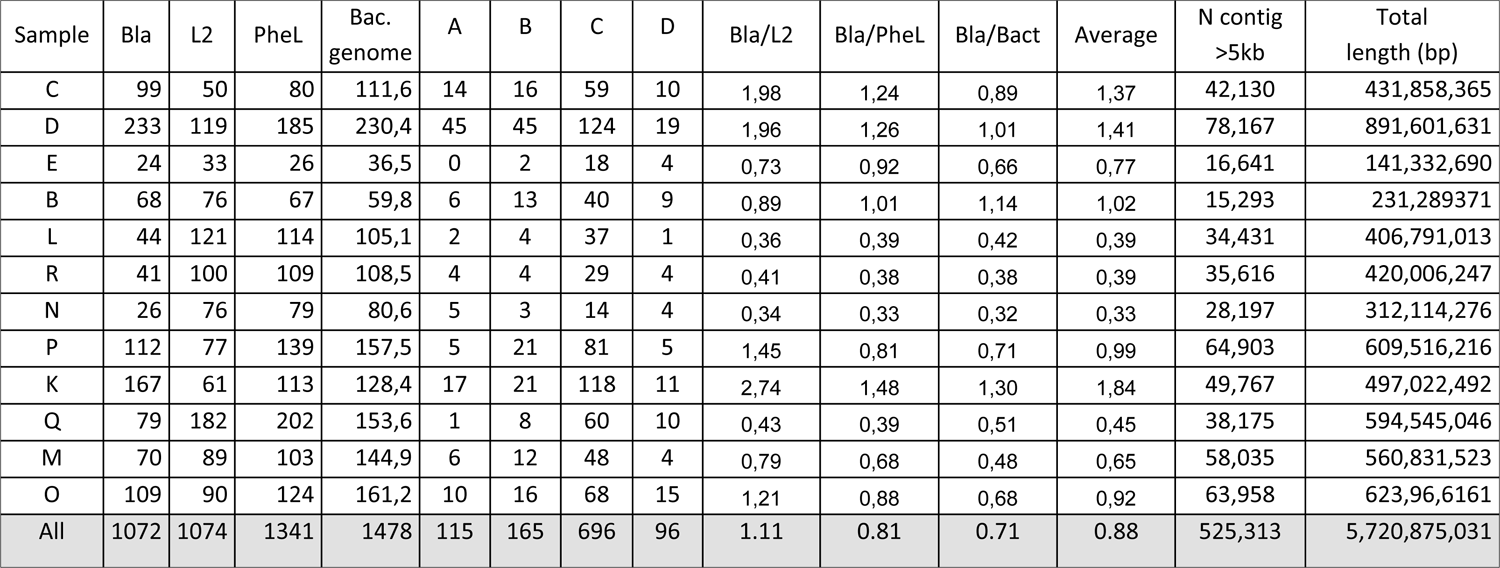
Analysis of contigs larger than 5 kb

| Sample | Bla | L2 | PheL | Bac.<br>genome | A | B | C | D | Bla/L2 | Bla/PheL | Bla/Bact | Average | N contig<br>>5kb | Total<br>length (bp) |
| --- | --- | --- | --- | --- | --- | --- | --- | --- | --- | --- | --- | --- | --- | --- |
| C | 99 | 50 | 80 | 111,6 | 14 | 16 | 59 | 10 | 1,98 | 1,24 | 0,89 | 1,37 | 42,130 | 431,858,365 |
| D | 233 | 119 | 185 | 230,4 | 45 | 45 | 124 | 19 | 1,96 | 1,26 | 1,01 | 1,41 | 78,167 | 891,601,631 |
| E | 24 | 33 | 26 | 36,5 | 0 | 2 | 18 | 4 | 0,73 | 0,92 | 0,66 | 0,77 | 16,641 | 141,332,690 |
| B | 68 | 76 | 67 | 59,8 | 6 | 13 | 40 | 9 | 0,89 | 1,01 | 1,14 | 1,02 | 15,293 | 231,289371 |
| L | 44 | 121 | 114 | 105,1 | 2 | 4 | 37 | 1 | 0,36 | 0,39 | 0,42 | 0,39 | 34,431 | 406,791,013 |
| R | 41 | 100 | 109 | 108,5 | 4 | 4 | 29 | 4 | 0,41 | 0,38 | 0,38 | 0,39 | 35,616 | 420,006,247 |
| N | 26 | 76 | 79 | 80,6 | 5 | 3 | 14 | 4 | 0,34 | 0,33 | 0,32 | 0,33 | 28,197 | 312,114,276 |
| P | 112 | 77 | 139 | 157,5 | 5 | 21 | 81 | 5 | 1,45 | 0,81 | 0,71 | 0,99 | 64,903 | 609,516,216 |
| K | 167 | 61 | 113 | 128,4 | 17 | 21 | 118 | 11 | 2,74 | 1,48 | 1,30 | 1,84 | 49,767 | 497,022,492 |
| Q | 79 | 182 | 202 | 153,6 | 1 | 8 | 60 | 10 | 0,43 | 0,39 | 0,51 | 0,45 | 38,175 | 594,545,046 |
| M | 70 | 89 | 103 | 144,9 | 6 | 12 | 48 | 4 | 0,79 | 0,68 | 0,48 | 0,65 | 58,035 | 560,831,523 |
| O | 109 | 90 | 124 | 161,2 | 10 | 16 | 68 | 15 | 1,21 | 0,88 | 0,68 | 0,92 | 63,958 | 623,96,6161 |
| All | 1072 | 1074 | 1341 | 1478 | 115 | 165 | 696 | 96 | 1.11 | 0.81 | 0.71 | 0.88 | 525,313 | 5,720,875,031 |

**Supplementary Table S2.** Analysis of 2 kb < contigs <5 kb

| Sample | Bla | L2 | PheL | Bact.<br>Genome | Bla/L2 | Bla/PheL | Bla/Bact | Average | N contig<br>2kb>L>5kb | Total bp |
| --- | --- | --- | --- | --- | --- | --- | --- | --- | --- | --- |
| C | 128 | 122 | 108 | 148,2 | 1,05 | 1,19 | 0,86 | 1,03 | 200,802 | 573,410,069 |
| D | 191 | 155 | 151 | 207,3 | 1,23 | 1,26 | 0,92 | 1,14 | 275,382 | 802,389,205 |
| E | 26 | 94 | 68 | 87,4 | 0,28 | 0,38 | 0,30 | 0,32 | 120,522 | 338,460,150 |
| B | 23 | 25 | 30 | 27,5 | 0,92 | 0,77 | 0,84 | 0,84 | 35,731 | 106,410,805 |
| L | 32 | 87 | 66 | 82,6 | 0,37 | 0,48 | 0,39 | 0,41 | 108,816 | 319,699,512 |
| R | 37 | 86 | 70 | 91,4 | 0,43 | 0,53 | 0,40 | 0,45 | 121,633 | 353,910,437 |
| N | 21 | 42 | 64 | 74,1 | 0,50 | 0,33 | 0,28 | 0,37 | 98,513 | 286,741,299 |
| P | 80 | 165 | 184 | 195,6 | 0,48 | 0,43 | 0,41 | 0,44 | 258,914 | 756,843,977 |
| K | 157 | 124 | 136 | 176,6 | 1,27 | 1,15 | 0,89 | 1,10 | 237,762 | 683,392,792 |
| Q | 33 | 69 | 83 | 85,4 | 0,48 | 0,40 | 0,39 | 0,42 | 112,563 | 330,646,859 |
| M | 69 | 133 | 157 | 181,4 | 0,52 | 0,44 | 0,38 | 0,45 | 241,064 | 701,920,384 |
| O | 125 | 141 | 150 | 181,4 | 0,89 | 0,83 | 0,69 | 0,80 | 239,974 | 701,853,105 |
| All | 922 | 1243 | 1267 | 1539 | 0.70 | 0.68 | 0.56 | 0.65 | 2,051,676 | 5,955,678,594 |

**Supplementary Table S3.** Processing of the datasets from Vishnivetskaya et al. (69)

| Accession | Source | Total bp after cleaning | N contig > 5kb | Average size<sup>1</sup> | Median size<sup>2</sup> | Median coverage<sup>2</sup> |
| --- | --- | --- | --- | --- | --- | --- |
| SRX2163490 | Bykovsky Peninsula | 18,845,162,025 | 4382 | 7758 | 1379 | 9.0 |
| SRX2163491 | Gydan Peninsula | 20,905,522,718 | 836 | 7509 | 1318 | 44. |
| SRX2163492 | Omolon River | 26,577,180,575 | 9136 | 6698 | 1395 | 6.95 |

1: computed on the contig > 5 kb ; 2: computed on all contigs>1 kb

